# Liver-like glycogen metabolism supports glycolysis in naked mole-rat heart during ischaemia

**DOI:** 10.1101/2024.06.09.598072

**Authors:** Amanda Bundgaard, Nini Wang, Iuliia Vyshkvorkina, Maria Sol Jacome Burbano, Maksym Cherevatenko, Theodoros Georgomanolis, Frederik Dethloff, Patrick Giavalisco, Jan-Wilm Lackmann, Gary R Lewin, Christian Frezza, Jane Reznick

## Abstract

As a subterranean eusocial mammal, the naked mole-rat faces a particularly challenging environment characterised by patchily available food, low O_2_ and high CO_2_ levels. In response, naked mole-rats have evolved a suite of molecular and physiological adaptations to survive extreme hypoxia. Yet, how naked mole-rats rewire their metabolism to protect the heart has not been comprehensively addressed. Here, we performed comparative analyses of naked mole-rat and mouse organs exposed to ischaemic conditions. We show that naked mole-rats have retained features of foetal cardiac metabolism replacing fatty acid utilisation for a unique type of carbohydrate metabolism largely dependent on glycogen. We found that naked mole-rats have co-opted specialised liver-like glycogen handling mechanisms in the heart. Amongst these is the expression of liver-specific enzyme isoforms and amylase, a digestive enzyme known for starch breakdown in saliva and intestine but whose biological role in glycogen processing has not been fully recognised. We show that amylase is rapidly activated in ischaemia and hydrolyses internal glycosidic bonds for more efficient downstream processing. This biochemical adaptation occurred in both mouse and naked mole-rat livers but exclusively in the naked mole-rat heart, which retained higher ATP levels by maintaining an increased glycolytic flux in an amylase-dependent mechanism. Overall, we discovered a previously unknown type of glycogen metabolism in the naked mole-rat that holds relevance to pathologies where glycogen plays a role. Furthermore, we describe a novel type of metabolic plasticity in the heart which may be harnessed for cardiac disease.

## Main

Glycogen, the primary storage form of glucose, is a rapid and accessible form of energy that can be supplied to cells upon an energetic demand. Glycogen is beneficial in ischaemic conditions and some anoxia-tolerant species like the freshwater turtle (*Chrysemys* and *Trachemys*) and crucian carp (*Carassius Carassius*) can survive for days in anoxic waters by using their abundant glycogen stores^1–3^. Three ATP molecules are produced for every glucose-6-phosphate (G6P) converted from glycogen versus 2 ATP from each glucose molecule, and this thriftier route to generate ATP may be one reason anoxia-tolerant animals evolved such dependence on glycogen^4^. The foetal heart develops in a low oxygen environment and also shows a high dependency on glycogen^5^. Accordingly, glycogen occupies 30% of the foetal cardiomyocyte cell volume in stark contrast to 2% of an adult cardiomyocyte^6^. Reduction in glycogen content is linked to a neonatal metabolic switch in cardiomyocytes, where fatty acid oxidation replaces anaerobic glycolysis and glucose oxidation in order to support the high adenosine triphosphatase (ATP) demands of an adult mammalian heart^7^. Due to pressures of its harsh subterranean environment^8,9^, the naked mole-rat (nmr) as a terrestrial mammal has evolved extraordinary resistance to extreme hypoxia^8,10,11^. In this study we focused on novel mechanisms that protect naked mole-rat hearts during ischaemia. We find extensive rewiring of naked mole-rat cardiac metabolism towards a foetal mode of energy generation and high glycogen storage. Furthermore, we uncovered unique forms of glycogen processing in the naked mole-rat heart which converge on liver-like mechanisms of glycogen metabolism including release of polysaccharides via a novel amylase-dependent mechanism. Such multi-faceted metabolic rewiring in an anoxia-resistant long-lived mammal reveals possibilities of metabolic plasticity of an adult heart which may be harnessed for understanding and treating human cardiac pathologies.

### Distinct metabolic response between naked mole-rat and mouse to ischaemia

To investigate the metabolic adaptations of the naked mole-rat and mouse, we performed comparative metabolomics analyses of heart and liver tissue at different durations of ischaemia (Fig. 1a). Principal component analysis (PCA) separated the metabolic profiles according to species and tissue (Fig. 1b). PCA on individual tissues revealed three distinct clusters in naked mole-rat tissues which corresponded to time spent in ischaemia and was therefore suggestive of two separate metabolic states defined by acute (5,10 minutes) and prolonged ischaemia (30,60 minutes). Furthermore, metabolic profiles in ischaemia clustered separately to baseline. (Fig. 1c and d). A similar pattern was observed in mouse, however there was not such strict separation between different timepoints in ischaemia (Fig. 1c and d).

**Fig. 1:**
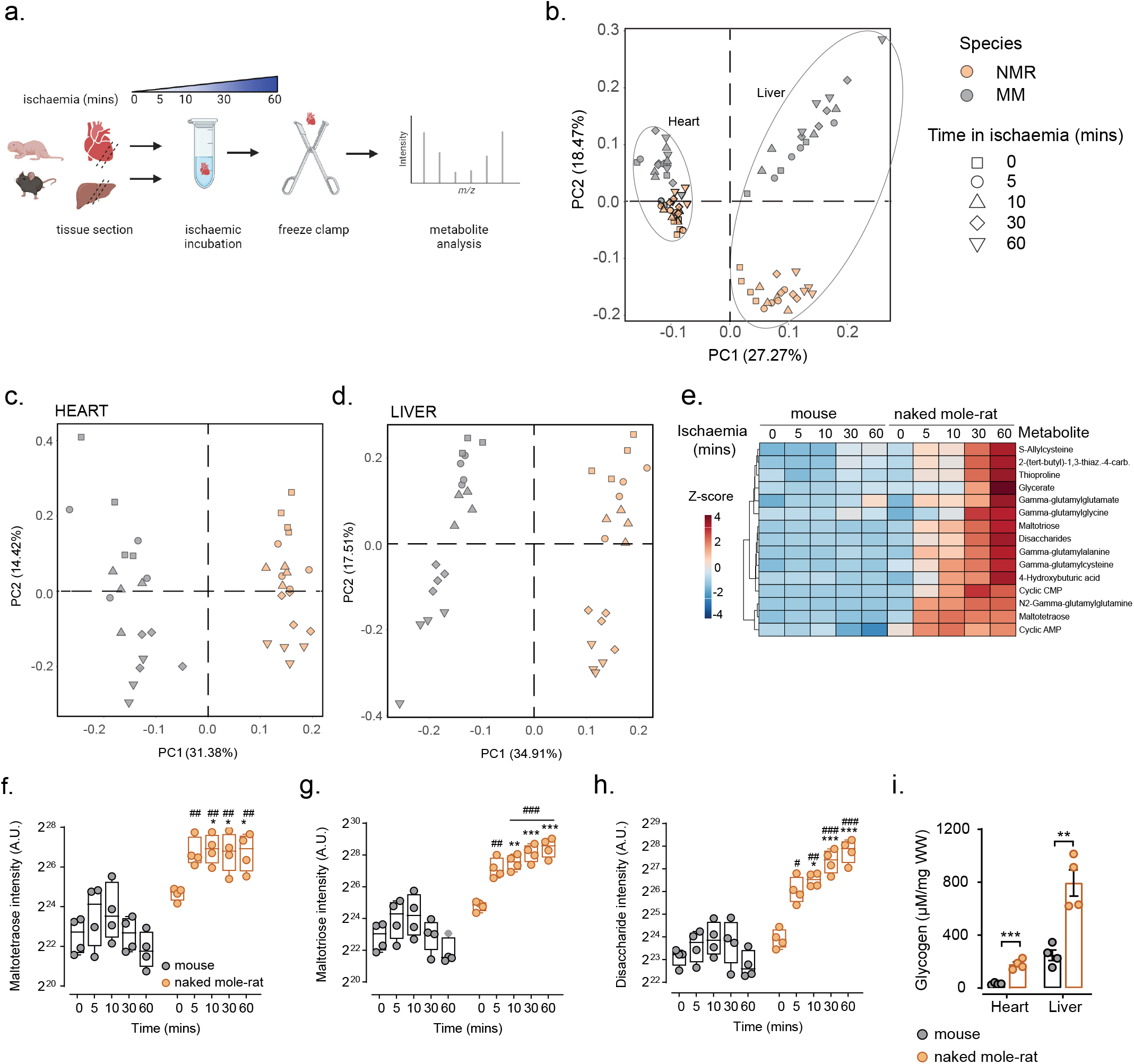
Distinct metabolic response to ischaemia in mouse and naked mole-rat heart. **a**, A schematic illustrating the experimental design to induce ischaemia in heart and liver sections in naked mole-rat and mouse. Sections from mouse and naked mole-rat heart and liver were either immediately clamped frozen at liquid nitrogen temperature to generate a baseline sample under fully oxygenated “normoxic” conditions or incubated to induce ischaemia at 30° for the indicated times before freeze clamping for downstream metabolomic analysis. **b-d**, Principal-component analysis of **b**, heart and liver **c**, heart and **d**, liver tissue from mouse and naked-mole-rats at baseline and exposed to ischaemia for 5,10,30 and 60 mins (n=4 each). **e**, Heatmap visualization of metabolites unchanged in mouse and naked mole-rat at baseline but increased in naked mole-rat ischaemic hearts (n=4 each). **f-h**, Levels of **f**, maltotetraose **g**, maltotriose **h**, and disaccharides in mouse and naked mole-rat hearts at different timepoints of ischaemia (n=4 each). **i**, Quantification of glycogen content in heart and liver in mouse and naked mole-rat (n=4). Error bar represents mean ± s.e.m. n numbers refer to individual animals. Two-way ANOVA with Tukey’s test was used for correction of multiple comparisons in **f-h**. Two-tailed, unpaired Student t-tests with correction for multiple testing were used for statistical analysis **i**. *p < 0.05, **p < 0.01, ***p < 0.001 within species comparison at different timepoints, #p < 0.05, ##p < 0.01, ###p < 0.001 between species comparison for corresponding timepoints.

### High glycogen content and release of polysaccharides in ischaemia

To identify metabolites which uniquely changed in naked mole-rat in response to ischaemia, but remained unchanged in mouse, we created a heatmap of all metabolites in heart or liver (Extended Data Fig 1a and b). Focusing on the heart, we narrowed down our analysis to metabolites that were the same in mouse and naked mole-rat in normoxia but increased in abundance in ischaemia in naked mole-rats only (Fig. 1e). Within the above cluster (Fig. 1e), disaccharide, maltotriose and maltotetraose appeared at similar levels between naked molerat and mouse in normoxia but were uniquely upregulated in naked mole-rat hearts across all ischaemic timepoints (Fig. 1f-h). Surprisingly, these three polysaccharides were similarly elevated in ischaemic liver in both species (Extended Fig. 1c-e) suggesting that naked molerat hearts have hijacked a liver-like response. The above polysaccharides may originate from a larger oligosaccharide, glycogen that represents the main storage of glucose in a cell and has been shown to be a crucial energy source during ischaemia and anoxia^6,12^. Consistent with previous findings^13,14^ we found a higher amount of glycogen in naked mole-rat hearts resembling levels found in liver, (Fig. 1i), the tissue with the highest glycogen storage capacity. Taken together, these data suggest a distinct metabolic response between naked mole-rat and mouse and a naked mole-rat-specific polysaccharide metabolism in the heart.

### Carbohydrate metabolism replaces fatty acid storage and use in naked mole-rat heart

We performed a transcriptomics analysis between naked mole-rat and mouse heart and livers to identify possible genes contributing to the metabolic phenotype in naked mole-rats. Using GO term enrichment analysis, we found that many genes related to the term Glycogen metabolic process were differentially expressed in hearts and livers between naked mole-rat and mouse (Fig. 2a, Extended Fig. 2a). Interestingly, carbohydrate and glycogen metabolism were amongst the top enriched GO terms for upregulated genes in naked mole-rat heart (Fig. 2b) and fatty acid and lipid metabolic process appeared as top downregulated GO terms (Fig. 2c). Lipid droplets (LD) are dynamic organelles storing neutral lipids for later use under energetic deficit^15–17^. LDs were readily detected with transmission electron microscopy (TEM) from sections of adult mouse heart but were completely absent in naked mole-rat heart sections. We then analysed LDs in neonatal mouse heart since the transcriptional profile favouring carbohydrate metabolism over fatty acids in naked mole-rats resembled a fetal-like cardiac programme^7^ and we supposed that lack of LDs may be a retained neotenous trait previously reported in naked mole-rat^18^. LDs were however present in neonatal mouse heart but at reduced numbers compared to adult mouse (Fig. 2d). Lack of LDs in naked mole-rat heart was further reflected by lower Oil Red O (ORO) staining which detects neutral lipids like triglyceride in tissue (Fig. 2e) and correlated with reduced expression of genes regulating cardiomyocyte lipid storage and lipid droplet dynamics^16,19^ (Fig. 2f). Genes in the GO term for fatty acid beta-oxidation, the primary mode of energy generation in adult heart were also downregulated in the naked mole-rat heart (Fig. 2g). Overall, this analysis suggests a major metabolic rewiring of heart metabolism in the naked mole-rat away from lipid utilisation and towards glucose storage and usage.

**Fig. 2:**
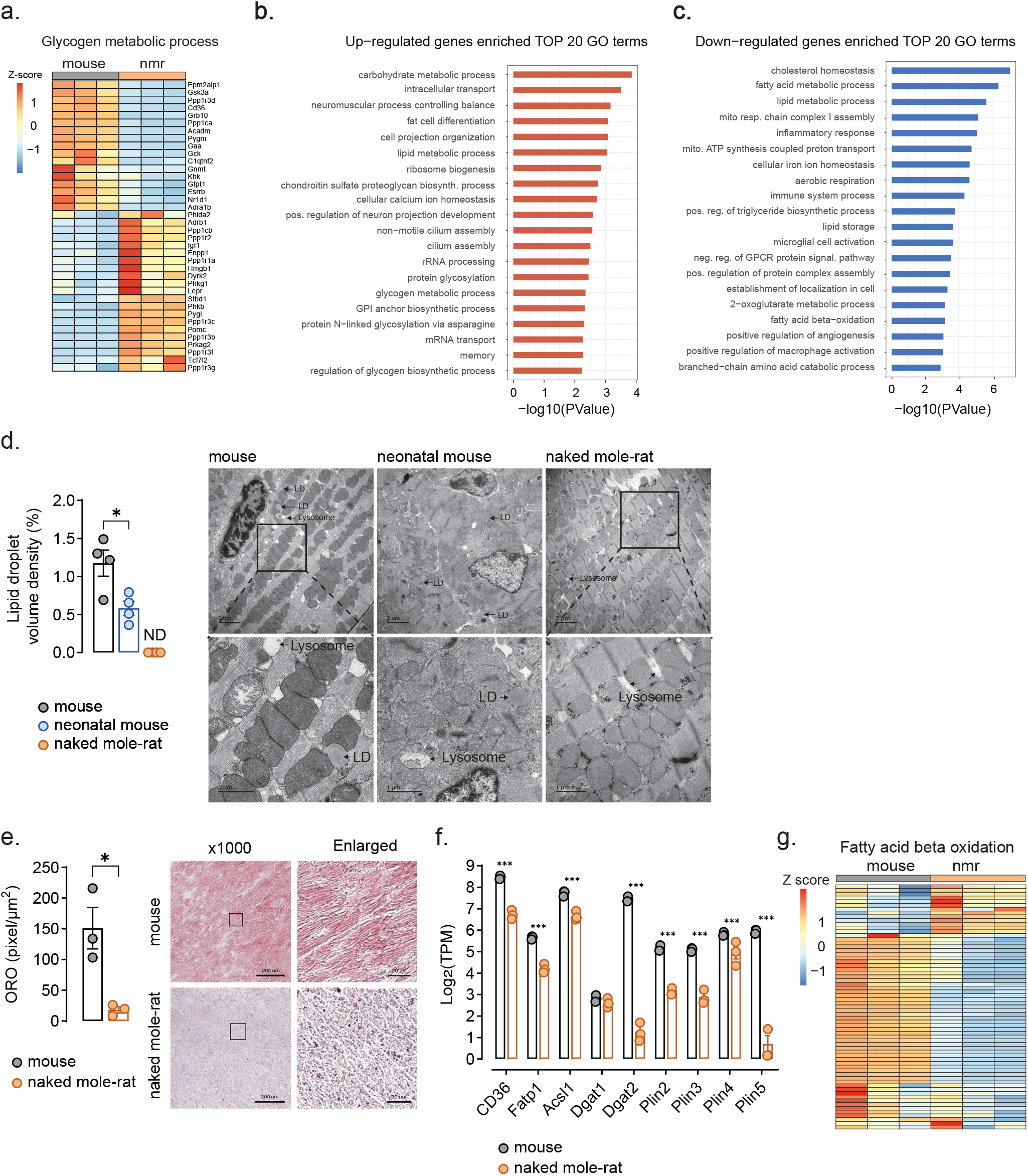
Carbohydrate metabolism replaces fatty acid storage and use in naked mole-rat heart. **a**, Heatmap of transcriptomic analysis between mouse and naked mole-rat heart for the GO term “glycogen metabolic process”. **b-c**, Top 20 enriched GO terms of b, upregulated genes and **c**, downregulated genes in naked mole-rat heart compared to mouse heart (n=3 for a-c). **d**, Quantification of % lipid droplet volume density in naked adult and neonatal (P1) mouse and adult naked mole-rat hearts and representative transmission electron microscopy images, scale bar = 2µM top panel, 1 µM bottom panel. Arrows point to representative lipid droplets (LD) and lysosomes (n=52-62 individual images for n=4 biological replicates). **e**,Representative Oil red O staining images indicating intramyocardial lipid content in mouse and naked mole-rat heart and quantitative analysis of positive area of oil red staining (30 randomly selected areas were measured in n=3 biological replicates). **f**, Expression of genes related to lipid droplet formation in heart of mouse and naked mole-rat determined with RNAseq (n=3). **g**, Heatmap of transcriptomic analysis between mouse and naked mole-rat heart for the GO term fatty acid beta oxidation. Error bar represents mean ± s.e.m. n numbers refer to individual animals, unless otherwise stated. Two-tailed, unpaired Student t-tests for **e**, with correction for multiple testing in **d, f** were used for statistical analysis, *p < 0.05, **p < 0.01, ***p < 0.001.

### Liver-like glycogen storage capacity in naked mole-rat heart

Glycogen is arranged in either α- or β-granules. β-granules are∼20–30 nm in diameter, consist of a central priming protein, glycogenin, covalently bound to a glucose polymer and are considered a rapid energy source. In contrast, α-granules are formed by several β-granules arranged in a broccoli-like fashion and are typically larger than 50nm and up to 300 nm in diameter^20^. α-Granules are mainly found in liver and have been linked to a slower release of energy^20^. Using transmission electron microscopy (TEM) imaging we could visualise glycogen granules within sections of adult mouse and naked mole-rat heart and liver and neonatal (P1) mouse heart (Fig. 3a). Correlated with high glycogen content (Fig.1i), naked mole-rat and neonatal mouse heart sections contained more electron-dense black glycogen granules compared to adult mouse, where glycogen granules appeared very rarely (Fig. 3a). Cardiac glycogen in adult and neonatal mouse consisted exclusively of β-granules smaller than 50 nm (Fig. 3a). Naked mole-rat heart however, contained high amounts of β – and surprisingly, a similarly high number of α-granules, mainly 80 nm but reaching up to 200-300 nm in diameter (Fig. 3a). Having observed α-granules in the naked mole-rat heart, we then compared naked mole-rat heart sections to livers of mouse and naked mole-rat where we observed as expected ample amounts of α-granules. Interestingly, glycogen granule size in naked mole-rat heart was not different to naked mole-rat liver where α-granules tended to be under 100nm and smaller than the α-granules observed in mouse liver (>100 nm) (Fig. 3a). Higher capacity of glycogen biosynthesis and storage in naked mole-rat was further reflected through almost 4- and 2-fold higher levels of glycogen precursor UDP-glucose in heart and liver, respectively (Fig. 3b and c). UDP-glucose dramatically dropped across ischaemic samples in naked mole-rat heart and liver and mouse liver (Fig. 3 b and c) suggesting a rapid switch towards glycogen breakdown under energy depleted states in these tissues. These data reveal that despite reliance on neonatal-like cardiac metabolism, naked mole-rats have diverged their glycogen from the neonatal form and evolved a unique way to store glucose in large liver-like granules.

**Fig. 3:**
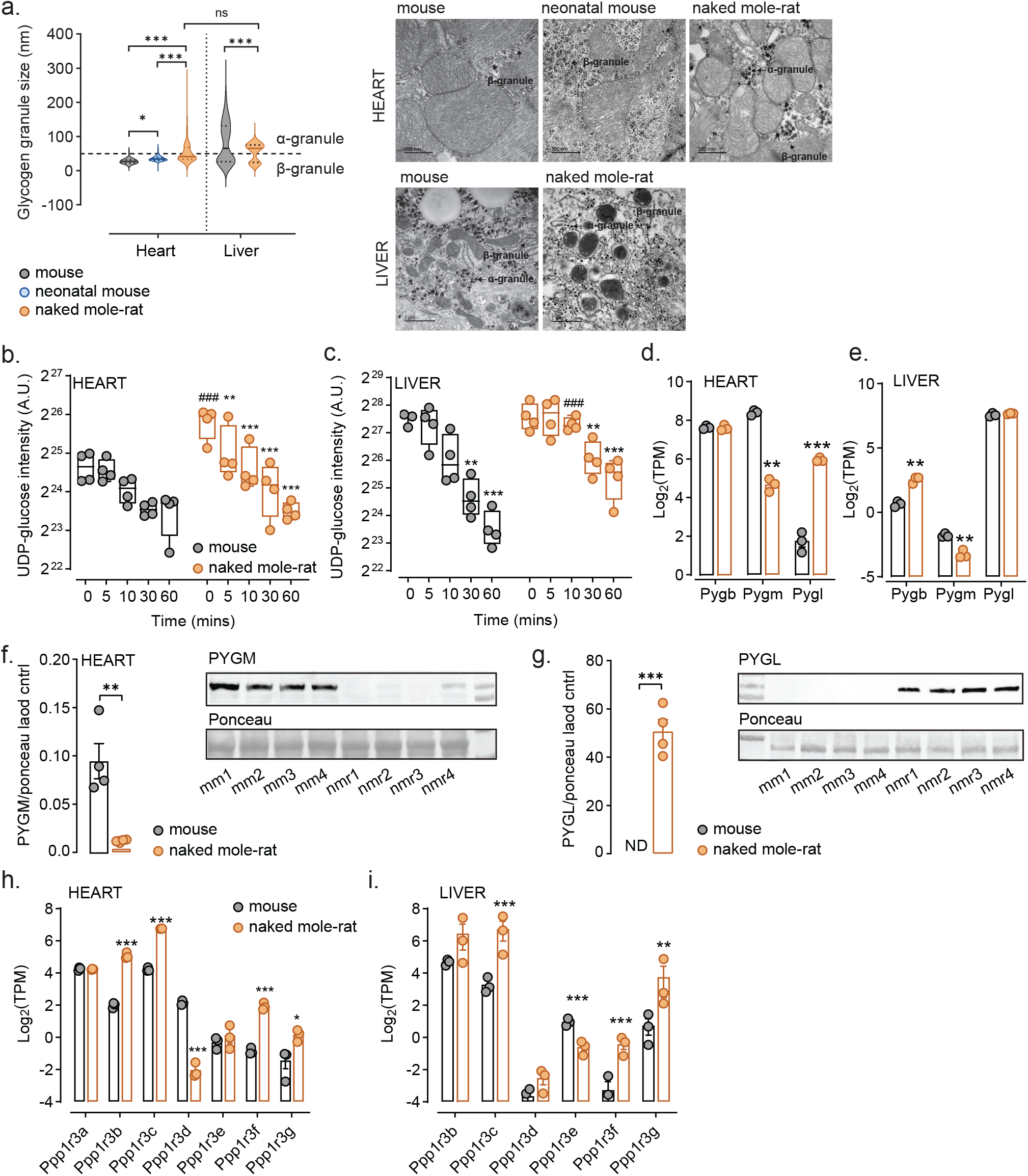
Glycogen storage and breakdown mechanisms resemble liver in naked mole-rat heart. **a**, Quantification of glycogen particle size and representative TEM images in adult and neonatal (P1) mouse and adult naked mole-rat heart (n=3), scale bar = 500nM and adult mouse and naked mole-rat liver (n=2), scale bar=1µM, a-granules < 50nm and -granules >50nm, (140-450 granules quantified for each sample). **b**,**c**, UDP-glucose levels at baseline and different lengths of ischaemia in **b**, heart and **c**, liver of mouse and naked mole-rat. **d**,**e**, Expression of glycogen phosphorylase (GP) isoforms Pygb, Pygm and Pygl in d, heart and **e**, liver determined by RNAseq (n=3). **f**,**g**, Western blot analysis and quantification normalised to ponceau loading control of GP isoforms in mouse and naked mole-rat heart (n=4) **f**, muscle isoform PYGM and **g**, liver isoform PYGL. **h**,**i**, Expression of Ppp1r3 isoforms in **h**, heart and **i**, liver determined with RNAseq (n=3). Error bar represents mean ±s.e.m. n numbers refer to individual animals. One-way ANOVA with Tukey’s test was used for correction of multiple comparisons in **a**, Two-way ANOVA with Tukey’s test was used for correction of multiple comparisons in **b, c**. Two-tailed, unpaired Student t-tests with correction for multiple testing were used for statistical analysis **d**,**e**,**h**,**i**. Two-tailed, unpaired Student t-tests were used for statistical analysis in f,g. *p < 0.05, **p < 0.01, ***p < 0.001, for b, c *p < 0.05, **p < 0.01, ***p < 0.001, within species comparison at different timepoints, #p < 0.05, ##p < 0.01, ###p < 0.001 between species comparison for corresponding timepoints.

### Naked mole-rat hearts express liver-specific glycogen handling isoforms

Amongst the differentially expressed genes related to glycogen metabolism (Fig2a), we observed preferences for non-canonical isoforms expressed in the heart. Particularly, the three isoforms of glycogen phosphorylase (GP) encoded by 3 genes *Pygb, Pygm and Pygl* designated brain, muscle, liver for the tissue where the respective isoform is predominantly expressed, showed altered pattern of expression. GP catalyzes the first step in glycogenolysis by releasing Glucose1-phosphate (G1P) from the terminal alpha-1,4-glycosidic bond of a glycogen molecule^21^. RNAseq analysis revealed that in mouse heart the two predominant isoforms are *Pygm* and *Pygb*, however in the naked mole-rat the isoform distribution is skewed to express substantially more *Pygl* at levels similar to the liver and much reduced *Pygm* expression (Fig. 3d and e). In naked mole-rat liver, isoform distribution of GP mimicked mouse liver (Fig. 3e). The isoform switch in the heart was even more pronounced at the protein level where we detected high levels of PYGM in mouse heart as expected and almost no protein expression in naked mole-rat heart (Fig. 3f). On the contrary, PYGL immunoblotting resulted in a strong signal in the naked mole-rat heart and there was no visible signal in the mouse (Fig. 3g). In liver, both species had almost undetectable level of PYGM protein expression and similarly high levels of PYGL protein between the two species (Extended Data Fig.2 b and c).

Similarly, genes in the *Ppp1r3* family, which control glycogen synthesis and breakdown, showed differential isoform preference in the naked mole-rat heart (Fig. 3h) with *Ppp1r3a*, the isoform canonically expressed in heart found at similar levels in mouse and naked mole-rat (Fig. 3h), but the liver isoforms *Ppp1r3b* and *Ppp1r3c* more abundantly expressed in naked mole-rat hearts (Fig. 3h). Of note, in the liver, *Ppp1r3b* did not differ in its mRNA expression but *Ppp1r3c* had elevated expression in naked mole-rats (Fig. 3i). Since high expression of *Ppp1r3b* and *Ppp1r3c* is linked to increased glycogen storage and glycogen granule size^22–24^ it is likely that alternative distribution of *Ppp1r3* isoforms and their overall increased expression in heart and liver is responsible for the enhanced capacity to store glycogen in naked mole-rat tissue. Overall, these results indicate that naked mole-rat hearts have non-canonical routes of glycogen storage and handling.

### Ischaemia promotes rapid breakdown of glycogen to maintain glycolytic flux

To understand the fate of glycogen under ischaemia we analysed glycogen content with a glycogen antibody^25^ immunostaining in heart sections before and after 30 minutes of ischaemia (Fig. 4a). There was a significant reduction in glycogen content in ischaemic conditions (Fig.4a) which we confirmed via glycogen quantification after 30 and 60 minutes of ischaemia (Fig. 4b)

**Fig. 4:**
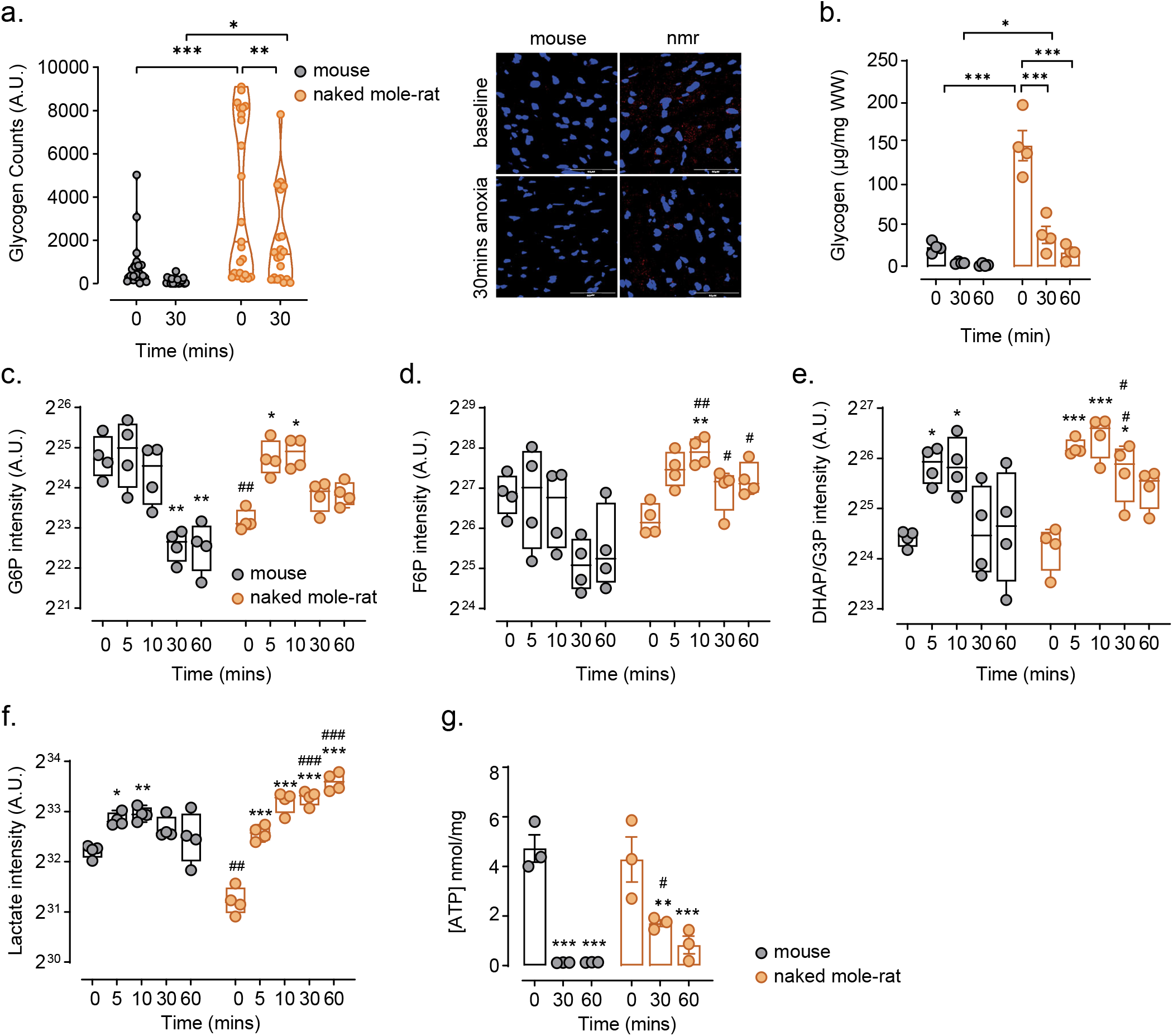
Naked mole-rats tap into glycogen stores to sustain glycolytic intermediates in anoxia. **a**, Counts of glycogen particles identified by immunofluorescence in heart tissue at baseline and after 30 min anoxia and representative immunofluorescent images of hearts. Blue: DAPI, red: glycogen, scale bar = 50μM (19-23 images were analysed for each species/condition, n=3 biological replicates). **b**, Quantification of heart glycogen content at baseline and 30 and 60mins ischaemia (n=4 each) in mouse and naked mole-rat (n=4). **c-f**, Levels of glycolytic intermediates at baseline and different duration of ischaemia in heart of mouse and naked mole-rat **c**, glucose-6-phosphate (G6) **d**, fructose-6-phosphate (F6P) **e**, DHAP/G3P and **f**, lactate (n=4 each). **g**, Quantification of ATP levels at baseline and 30 and 60 mins of ischaemia in mouse and naked mole-rat heart (n=3). Error bar represents mean ± s.e.m. n numbers refer to individual animals in **d-i**. Two-way ANOVA with Tukey’s test was used for correction of multiple comparisons in **a-g**. *p < 0.05, **p< 0.01, ***p < 0.001, for **c-g**, *p < 0.05, **p < 0.01, ***p < 0.001, within species comparison at different timepoints, #p < 0.05, ##p < 0.01, ###p<0.001 between species comparison for corresponding timepoints.

Synthesis and breakdown of glycogen referred to as the “glycogen shunt” is channelling glucose via glycogen to produce Glucose-6-phosphate (G6P), which ensures homeostasis of metabolic intermediates and maintenance of cellular energy in the form of ATP through local and thus rapid access to glucose^26^. Hence, we analysed intermediates of upper and lower glycolysis including G6P, F6P, DHAP/3PGA, and lactate from our *ex vivo* ischaemic heart metabolomics dataset (Fig. 4c-f, Extended Data Fig. 3a and b). Metabolites of upper glycolysis as well as DHAP/3-PGA rapidly increased in the early phase in naked mole-rat heart (5 and 10 mins) and then declined in prolonged ischaemia (30 and 60 mins) but remained significantly higher than baseline suggestive of a new steady-state. In contrast, in mouse heart, despite an initial surge of these metabolites, glycolytic intermediates could not be maintained at longer duration of ischaemia and significantly decreased below baseline. Furthermore, lactate continued to incrementally increase in naked mole-rat heart the longer the tissues remained ischaemic (Fig. 4f) whereas in mouse, lactate was only marginally increased at early timepoints of ischaemia and stagnated at 30 and 60 minutes. Since this experiment was carried out on excised tissue cut off from circulation and not submerged in buffer, lactate could not be exported from the tissue and thus served as a proxy for glycolytic flux which continued running in naked mole-rats throughout 60 minutes of ischaemic exposure but halted between 10-30 minutes in mouse (Fig. 4f). As expected, ATP levels dramatically dropped in ischaemia in both species, but correlating with higher levels of glycolytic intermediates, ATP was nevertheless maintained at 10-fold higher levels in naked mole-rat compared to mouse at 30 minutes of ischaemia (Fig. 4g).

### Amylase is activated in ischaemia to process glycogen into polysaccharides

Considering the well characterised canonical glycogen breakdown pathway (Fig.5a), the appearance of polysaccharides in ischaemia was surprising and prompted us to look for other non-canonical means of processing glycogen. α-Amylase is a digestive enzyme that catalyzes the hydrolysis of internal α-1,4-glycosidic bonds of starch into smaller polysaccharides such as maltose, maltotriose and maltotetraose^27^. Mammalian α-amylase is mainly synthesized by the pancreas and salivary glands but there have been reports of much lower amounts of α-amylase mRNA and activity detected in rat liver ^28–30^, intestine ^31^, brain ^32,33^ and other tissues^34^. The biological significance of α-amylase outside of salivary gland and pancreas remains unclear.

Since α-amylase has the capacity to break down glycogen molecules, we decided to investigate whether polysaccharides detected in heart and liver in ischaemia are products of α-amylase activity. We scrutinised our transcriptomics data for amylase expression in heart and liver tissue and found low expression of amylase in mouse liver which was nevertheless over ten-fold higher than heart tissue (Fig. 5b and c, Extended Data Fig. 4a and b) confirming previous reports that liver indeed expresses amylase^28,29,35^. Naked mole-rat liver showed 6-fold higher expression than mouse. Most remarkably, we detected significant expression of amylase in heart tissue, albeit relatively low compared to liver (Fig. 5b, Extended Data Fig. 4a and b). The transcript reads mapped to *amy1* gene in both heart and liver in mouse and naked mole-rat (Extended Data Fig. 4a and b). Therefore, amylase found in tissues outside of pancreas and salivary gland is *Amy1*, corroborating previous studies reporting liver amylase expression to originate from the amy1 gene^28^. To check that the mRNA transcript was converted into functional protein we performed western blot analysis and an amylase activity assay. Amylase has been reported to be between 56-62 kD depending on its glycosylation state^36^. We were able to detect amylase in the livers of both mouse and naked mole-rat, with the band for naked mole-rat amylase showing a slightly slower migration suggesting a more glycosylated state (Fig. 5e). Quantification of the bands indicated that naked mole-rats express 2.5-fold more amylase protein in liver. Moreover, we detected a clear band for amylase at about 60kD in heart tissue, but only in the naked mole-rat (Fig. 5d). Mass spectrometry-based protein analysis confirmed the presence of amylase in mouse and at greater intensity naked mole-rat heart and liver as well as naked mole-rat brain (Extended Data Fig. 4c). Finally, an activity assay on lysates from heart and liver showed that amylase activity in naked mole-rat heart was over 6-fold higher than mouse and similar to levels in the mouse liver. We were also able to substantially inhibit the activity of amylase with the application of an amylase inhibitor, acarbose (Fig. 5f). Interestingly, mouse heart exhibited some amylase activity albeit at very low levels which was nevertheless reduced to non-detectable activity with acarbose suggesting functional amylase in mouse heart. Overall, elevated expression and activity of amylase in heart of the naked mole-rat may be an evolved adaptation to gain faster access to energy in hypoxia.

**Fig. 5:**
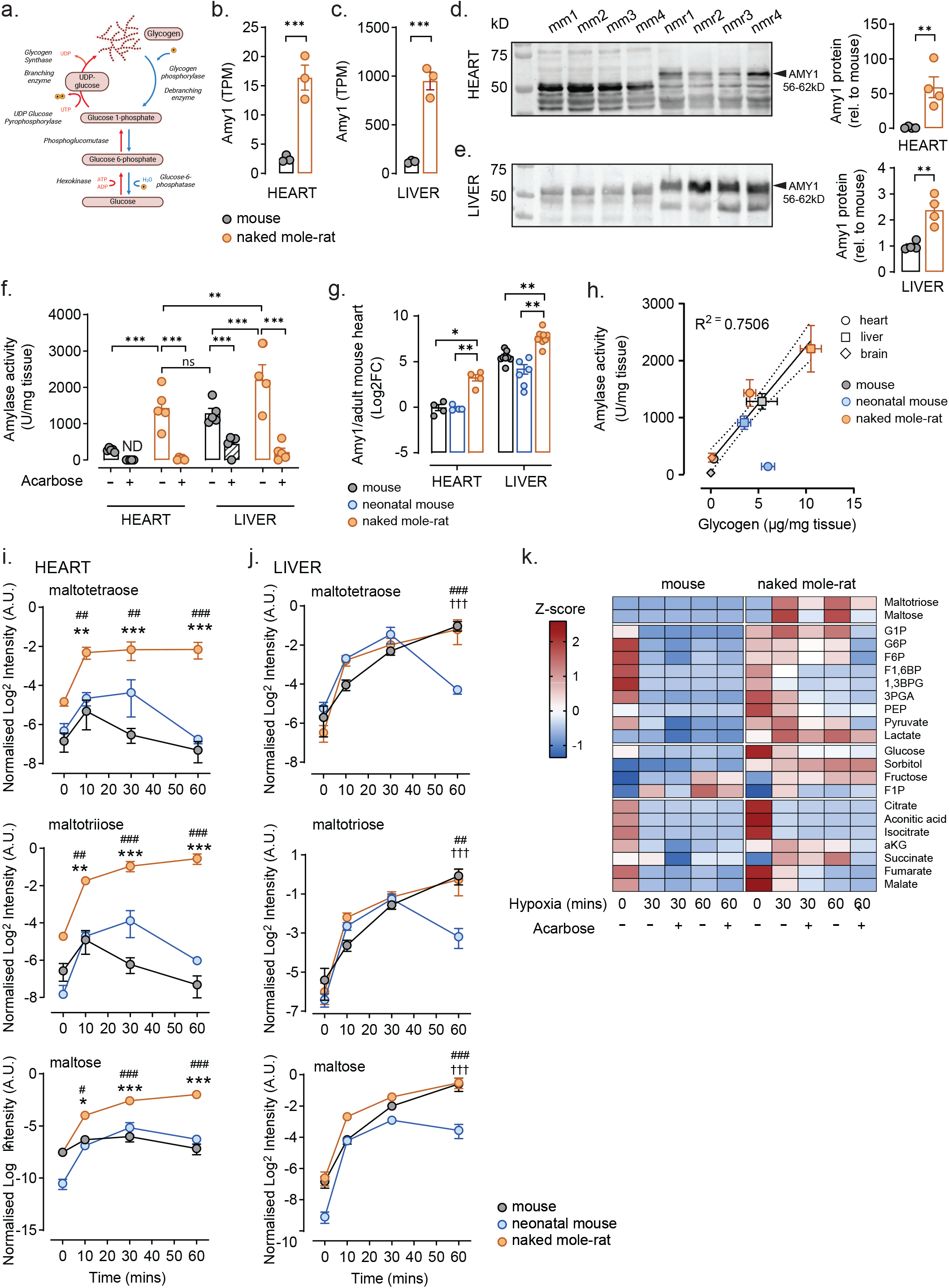
Naked mole-rat has co-opted amylase in the heart for efficient glycogen breakdown. **a**, Canonical glycogen synthesis and breakdown pathways. **b**,**c**, Expression of salivary amylase (Amy1) in b, heart and c, liver determined by RNAseq (n=3). **d**,**e**, Western blot analysis of amylase protein in **d**, heart and **e**, liver of mouse and naked mole-rat and quantification of amylase protein relative to mouse (n=4 each). AMY1 predicted size is 56-62 kDa depending on glycosylation status for validation of bands with LC/MS e, Amylase activity in tissue homogenates in heart and liver with and without amylase inhibitor acarbose (2.5mM) (n=5, except nmr liver n=4). **f**, Expression of Amy1 in adult mouse, naked mole-rat heart and liver and neonatal (P1) mouse heart relative to adult mouse heart measured by qPCR (heart n=4, neonatal mouse liver n=6, mouse and nmr liver n=9). **g**, correlation between amylase activity and glycogen content in heart, liver and brain tissue from adult and neonatal (P1) mouse and adult naked mole-rat estimated by simple linear regression and 95% confidence interval, R2=0.7506, p<0.0001 (n=5, except nmr liver, brain and mouse brain n=4). **h**,**i**, Intensity (normalised to median of L-Valine-d8) of maltotetraose, maltotriose and maltose in adult, neonatal mouse and naked mole-rat at normoxic baseline and indicated timepoints in ischaemia in h, heart and **i**, liver (n=4 each). **j**, Heatmap of all measured metabolites in mouse and naked mole-rat heart slices in 0.1% hypoxia with or without acarbose treatment. Intensities were normalised to corresponding hypoxia 30 min sample, n=4. Error bar represents mean ± s.e.m. n numbers refer to individual animals. Two-tailed, unpaired Student t-tests with correction for multiple testing were used for statistical analysis in **b**,**c**,**g**. Two-tailed, unpaired Student t-tests were used for statistical analysis in d,e. Two-way ANOVA with Tukey’s tests was used for correction of multiple comparisons in f, h-j. *p < 0.05, **p < 0.01, ***p < 0.001, for **h** and **i**, * indicates significance between naked mole-rat vs adult mouse, # naked mole-rat and neonatal mouse, † adult and neonatal mouse.

Neonatal hearts contain several-fold more glycogen than adult hearts (Fig. 3a) and we questioned whether glycogen and therefore amylase in the heart may be a retained neonatal feature given that naked mole-rats indeed have several molecular remnants of neoteny^18,37^ including a glucose dependent metabolism in the heart (Fig. 2). We analysed amylase expression by qPCR but found the expression of amylase in neonatal heart at similar level to adult mouse heart (Fig. 5g) suggesting that elevated cardiac amylase expression is a uniquely naked mole-rat feature. To confirm a functional link between amylase and glycogen, we measured glycogen levels in three different adult tissues (brain, heart and liver) and neonatal mouse heart and liver tissue and correlated it to amylase activity for the corresponding organ. Except for neonatal heart, there was a remarkable correlation across all tissues analysed where the glycogen content very closely correlated with the tissue’s amylase activity (R^2^=0.7506) (Fig. 5h). Linear regression analysis revealed that neonatal mouse falls outside of the 95% confidence interval due to amylase activity being very low relative to the high glycogen content of the neonatal heart) (Fig. 5h). Finally, we exposed neonatal mouse heart and liver to different times of ischaemia and compared levels of maltotetraose, maltotriose and maltose with data from adult mouse and naked mole-rat (Fig. 5i and j). In the heart, all three polysaccharides were significantly more abundant in the naked mole-rat across all timepoints compared to both adult and neonatal mouse (Fig. 5i) correlating with much higher amylase activity in naked mole-rat heart (Fig. 5h). Interestingly, both adult and neonatal mouse showed some increase in polysaccharides in ischaemic conditions where neonates maintained elevated polysaccharide levels for a longer duration than adult mice (30 mins versus 10 mins ischaemia) (Fig. 5i). This suggests that there is indeed low amylase activity induced by ischaemia in adult and neonatal mouse heart and it is likely that, since the neonatal mouse has more glycogen as substrate for its amylase, correspondingly, glycogen is broken down to polysaccharides for a longer duration in neonates in ischaemia. All three animal groups increased polysaccharide levels dramatically in ischaemic liver (Fig. 5j) correlating with our observations of high amylase expression and activity in liver of both species (Fig. 5h). This data supports a biological role for amylase in ex-pancreatic and ex-salivary tissue and reveals an unrecognised role for amylase activity in heart and liver to be rapidly stimulated in ischaemia to release polysaccharides from glycogen.

### Inhibition of amylase results in reduction of glycolytic intermediates and F1P

To understand the biological significance of glycogen breakdown via amylase under low oxygen conditions, we performed an *ex vivo* experiment where we exposed small tissue pieces of heart to 30 and 60 mins of extreme hypoxia (0.1% O2) with and without the amylase inhibitor acarbose followed by mass spectrometry-based metabolite analysis. A scheme of the metabolites measured is depicted (Extended Data Fig. 5a) and metabolites that were significantly reduced with acarbose in the naked mole-rat and mouse are coloured in blue. The heatmap in Fig. 5j reveals that many glycolytic metabolites are present at similar levels in naked mole-rat and mouse heart at baseline normoxic conditions. However, in extreme hypoxia the mouse rapidly and dramatically drops its levels of glycolytic intermediates whereas the naked mole-rat can maintain glycolytic metabolites at steady levels up to 30 minutes and for some metabolites, even up to 60 minutes (Fig. 5j, Extended Data Fig. 5a, f-l), re-affirming our previous observation that naked mole-rats can sustain glycolytic intermediates for much longer under extreme energetic challenges. Inhibition of amylase via acarbose results in a significant reduction of glycolytic, fructolytic and some TCA cycle intermediates implicating amylase in facilitating efficient glycolytic flux in hypoxia. Supporting our initial observation (Fig. 1e-h), we saw a dramatic increase of maltotriose and maltose in the naked mole-rat heart upon extreme hypoxia which was severely blunted by the addition of acarbose (Fig. 5j, Extended Data Fig.5 b and c). Compared to naked mole-rat hypoxic heart, polysaccharide content in mouse was magnitudes lower (Extended Data Fig.5 b and c), most likely since 30 minutes of extreme hypoxia is enough time to deplete the entire store of glycogen in mouse heart (Fig 4a and b). Glycogen and polysaccharides can be processed into G1P by GPs or into glucose by the action of α-glucosidase, GAA in the lysosome^38^ or MGAM^39^, a digestive enzyme like amylase, usually associated with processing polysaccharides into free glucose in the intestine (Extended Data Fig. 5a). G1P increased in hypoxia in naked mole-rats with a tendency to decrease with amylase inhibition but did not reach significance. In contrast, G1P levels in mice plummeted with hypoxia and were over 10-fold lower compared to naked molerat (Extended Data Fig. 5 e). Glucose was maintained over 4-fold higher in hypoxic naked mole-rat heart compared to mouse but decreased with acarbose treatment (Extended Data Fig. 5d). Surprisingly, sugar phosphates of the upper glycolysis G6P and F6P were unaffected by acarbose treatment (Extended Data Fig. f and g), in contrast to F1,6BP, a product of the rate-limiting PFK1 enzyme as well as all other downstream glycolytic intermediates (1,3BPG, 3PGA, pyruvate, lactate) which were drastically reduced in the presence of acarbose (Extended Data Fig. 5h-l). Acarbose showed similar effects on glycolytic intermediates in mouse (Extended Data Fig. 5a and h-l) which shows that mice rely somewhat on amylase, however due to overall low glycogen levels and much lower amylase expression and activity, amylase can only enhance glycolysis for a short interval until glycogen is depleted (Fig. 4a and b). Acarbose decreased TCA cycle intermediates in both species but increased succinate/fumarate ratio in naked mole-rats suggesting a more rapid inefficiency of mitochondria when amylase is blocked (Extended Data Fig. 5m-t).

We have previously reported that naked mole-rats switch to fructose metabolism during ischaemic episodes in liver, kidney and brain^10^, and here we show that the heart similarly raises its levels of fructose and F1P levels in extreme hypoxia (Fig.5k, Extended Data Fig. 5v and w). Surprisingly, a similar pattern was observed in mice (Fig.5k, Extended Data Fig. 5v and w). Since neither fructose nor glucose was provided in the media, the fructose and downstream F1P had to be generated *de novo* via the polyol pathway^40^ from free glucose. Free glucose can be generated from glycogen by non-canonical glycogen mobilization pathways via either α-glycosidases MGAM whose mRNA expression was 4-fold higher in naked mole-rat compared to mouse heart (Extended Data Fig. 6a) or GAA which we showed through immunostaining to be expressed at greater levels and have higher co-localisation with glycogen in the naked mole-rat heart (Extended Data Fig. 6b-d). Although neither fructose nor sorbitol was changed with acarbose treatment (Extended Data Fig. 5u and v), likely due to dynamic synthesis and breakdown fluxes, we nevertheless observed a significant reduction in F1P in hypoxia with acarbose treatment in both species (Extended Data Fig. 5w). Glycogen, therefore, may not only provide carbons for the classic glycolytic pathway, but may also act as a reservoir of free glucose released by α-glucosidases. Free glucose can be converted to fructose and F1P allowing bypass of the tightly regulated rate-limiting PFK1 enzyme to enter glycolysis downstream^9^. Indeed, it has now been shown in several systems that fructose metabolism is upregulated and beneficial under low oxygen conditions^9,41–43^. Overall, in both mice and more pronounced in the naked mole-rat where amylase plays a greater role, inhibition of glycogen mobilisation results in a dramatic reduction of glycolytic flux and therefore energetic resources which are vital, particularly under energy depleted conditions.

## Discussion

Naked mole-rats are amongst the most impressive hypoxia tolerant mammals, surviving 18 minutes of complete anoxia and hours at 3% O_2_^10,44^. We sought to understand mechanisms that afford naked mole-rat hearts protection under ischaemic conditions and uncover a dramatic rewiring of cardiac metabolism which forgoes lipid storage and utilisation in substitution for abundant glycogen stores and carbohydrate metabolism. Such metabolic rewiring is reminiscent of a foetal heart which because of its primitive tubular morphology and intrauterine environment is inherently hypoxic^45^. However, foetal metabolism cannot meet the energetic demand of a post-natal heart and soon after birth mammalian heart reprogrammes to a more efficient energy production using fatty acid oxidation^46^. To overcome energetic crises and sustain cardiac output in adulthood, we discovered that naked mole-rats optimised the foetal mode of energy generation via several unique adaptations resembling liver-like glycogen metabolism. This includes larger and more abundant glycogen granules arranged in α-particles normally found exclusively in liver^20^ and likely a result of increased expression of liver isoform PPP1R3B and PPP1R3C, previously linked with increased glycogen content^22–24^ and larger granule size^47^ respectively. Interestingly, larger α-particles release glucose more slowly than β-particles^48^, and help with maintaining blood glucose during overnight fasting^24^. In naked mole-rats α-granules may provide stable reservoirs of glucose for baseline cardiac function in the absence of lipid metabolism and simultaneously be an abundant local source of anaerobic fuel for energetic crises like ischaemia. Liver (PYGL) and muscle (PYGM) glycogen phosphorylases are both activated by phosphorylation, but in the unphosphorylated (b) state, only PYGM is efficiently activated by the allosteric activator AMP^49,50^. Additionally, PYGL is overall a less efficient enzyme than PYGM and altogether these differences reflect the distinct physiological roles for these two isoenzymes^50^. The liver stores by far the largest amount of glycogen compared to all other tissues and liver’s specialised role as a “glucostat” for systemic glucose homeostasis diversified its glycogen metabolism away from other tissues like muscle and brain where glycogen is primarily used to meet intracellular or intra-organ energy demands^51^. Replacement of PYGM for the liver PYGL isoform in naked mole-rat heart favours a hormonal control of glycogen breakdown dictated by whole-body energy status possibly avoiding a premature switch for glycogenolysis during transient intracellular energetic fluxes signalled via AMP. Likewise, lower specific activity of PYGL may protect naked molerats from inappropriately high rates of glycogen breakdown needed during fight-or-flight response in muscle but would be wasteful under baseline states or extended periods of hypoxia where energetic demands are lowered and substrate spared.

Under severe energetic challenges like ischaemia, we report an unexpected role for amylase enzyme to hydrolyse glycogen into smaller polysaccharides in heart and liver. Amylase is classically known to be expressed in the pancreas (*Amy2*) and salivary gland (*Amy1*) for starch breakdown ^27^ as well as liver^28^ where the biological function remained so far obscure. Naked mole-rats evolved higher expression of *amy1* gene compared to mouse in liver and uniquely in the heart. Amylase hydrolyses internal glycosidic bonds to yield short polysaccharides^52^ and in this way provides greater substrate for downstream processing by glycogen phosphorylase and glucosidases. Indeed we show amylase to be important for maintaining efficient glycolytic flux, adequate levels of free glucose and fructolysis in near-anoxic conditions. We mined RNAseq and metabolomics data from a recent study on cardiometabolic adaptations in 8 African mole-rats species^14^ and found that rewiring of glycogen metabolism including amylase expression in the heart occurred exclusively in the naked mole-rat genera and may indeed be a significant factor contributing to this species’ superior survival in extreme hypoxia^10,11^.

Glycogen has recently been implicated in many processes outside of energy storage including fibrosis, gene regulation and protein glycosylation in various tissues^53–55^. We believe the alternative way the naked mole-rat uses its glycogen stores could contribute insights to these recent developments. Moreover, the novel role for amylase in glycogen breakdown may not only offer insights into protective mechanisms during energy deficits like ischaemia, but may be a novel avenue to explore in dysregulation of glycogen metabolism in diseases like diabetes.

## Acknowledgements

We would like thank Paul Friedemann Pohlig and Mandy Götsche of the CECAD in vivo Research Facility for naked mole-rat colony management and care, CECAD Imaging Facility for microscopy support and the Regional Computing Center of the University of Cologne (RRZK) for providing computing time on the DFG-funded (Funding number: INST 216/512/1FUGG) High Performance Computing (HPC) system CHEOPS as well as support. We thank Michael P. Murphy and Andrew M. James for valuable inputs and discussion during the initial development of this project and Nick Adams for glycogen granule analysis. This work was supported by the Deutsche Forschungsgemeinschaft (DFG, German Research Foundation) under Germany’s Excellence Strategy - EXC 2030 – 390661388, the ERC Starting Grant to J.R. (grant ID 851653), a Villum International Postdoc Fellowship to A.B. (grant number 34435), a large instrument grant INST 216/1163-1 FUGG by the German Research Foundation (Großgeräteantrag der Deutschen Forschungsgemeinschaft).

## Author Contributions

A.B., and J.R. conceived the study and designed the experiments. A.B. performed most experiments with support of I.V. and M.S.J.B. Resources provided by G.R.L. N.W. and T.D. performed bioinformatics analysis. M.C., C.F. F.D. and P.G. performed and analysed metabolomics data. J.W.L. performed and analysed proteomics data. J.R. wrote the manuscript with input from all authors.

