## Extended Data Figure 1-6 for "Liver-like glycogen metabolism supports glycolysis in naked mole-rat heart during ischaemia"

**Extended Data Fig. 1: Metabolic response in ischaemic mouse and naked mole-rat heart and liver**

**a,b** Heatmaps of all metabolites detected in **a,** heart and **b,** liver from mouse and naked mole-rats in 0, 5, 10, 30 and 60 min of ischaemia. Black box in **a,** outlines the cluster represented in Fig. 1e.**c-e,** Levels of **c,** maltotetraose **d,** maltotriose and **e,** disaccharide in liver from mouse and naked mole-rats (n=3 for mouse liver at 10 mins, n=4 each for other groups for **a-e**). Error bar represents mean ± s.e.m. n numbers refer to individual animals. Two-way ANOVA with Tukey's test was used for correction of multiple comparisons in **c-e**. *$p < 0.05$, **$p < 0.01$, ***$p < 0.001$ within species comparison at different timepoints.

**Extended Data Fig. 2: Expression of glycogen related genes in mouse and naked mole-rat liver**

**a,** Heatmap of transcriptomic analysis between mouse and naked mole-rat heart and liver for the GO term glycogen metabolic process. **b,c**, Western blot analysis and quantification normalised to ponceau loading control of GP isoforms in mouse and naked mole-rat liver (n=4) **b,** muscle isoform PYGM and **c,** liver isoform PYGL (for gel source data see Supplementary Data 5). (n=4 each). Error bar represents mean ± s.e.m. n numbers refer to individual animals, unless otherwise stated. Two-tailed, unpaired Student t-tests were used for statistical analysis in **b,c**.

**Extended Data Fig. 3: Glycolytic intermediates in ischaemic heart of mouse and naked mole-rat**

Levels of glycolytic intermediates **a,** 2/3-phosphoglyceraldehyde (2/3PGA) and **b,** phosphopenolpyruvate (PEP) in baseline and ischaemia in mouse and naked mole-rat hearts (n=4 each). Error bar represents mean ± s.e.m. n numbers refer to individual animals. Two-way ANOVA with Tukey's test was used for correction of multiple comparisons. *$p < 0.05$, **$p < 0.01$, ***$p < 0.001$, within species comparison at different timepoints, $^{\#}p < 0.05$, $^{\#\#}p < 0.01$, $^{\#\#\#}p < 0.001$ between species comparison for corresponding timepoints.

**Extended Data Fig 4: Co-option of Amy1 in naked mole-rat heart for efficient glycogen breakdown**

**a,b,** Mapped reads for amylase from RNAseq analysis onto the **a,** mouse and **b,** naked mole-rat genomes. **a,** Mouse has 7 copies of amylase genes and **b,** naked mole-rat has 3 copies of amylase genes. Amylase reads from heart and liver map to Amy1 gene in both naked mole and mouse. Intensity of mapped reads is much higher in naked mole-rat for corresponding mouse tissue (n=3). **c**, in-gel proteomics analysis of selected size intervals cut from gel of proteins from mouse and naked mole-rat heart, liver and brain tissues separated by SDS-PAGE. Numbers indicate peak intensity (log2) of amylase protein found in corresponding gel piece with size indicated on the left, ND denotes that amylase was not detected (n=1).

**Extended Data Fig. 5: Inhibition of amylase decreases glycolytic and TCA cycle intermediates in hypoxia**

**a**, Summary of glycogen breakdown pathways in naked mole-rat and mouse heart, effect of acarbose on amylase in indicated. Metabolites that decrease with amylase inhibition in 0.1% hypoxia compared to hypoxia control are marked in blue, increased in yellow, unchanged metabolites in grey. Relative changes in levels normalized to 30 min hypoxia in naked mole-rat and mouse heart of **b,c** the polysaccharides **b,** maltotriose, and **c,** maltose **d**, glucose, **e,** glucose-1-phosphate (G1P) . Intermediates of upper glycolysis **f,** Glucose-6-phosphate, **g,** fructose-6-phosphate. Intermediates of lower glycolysis **h,** fructose-1,6-bisphosphate (F1,6BP) **i,** 1,3-bisphosphoglycerate (1,3BPG) **j,** 3-phosphoglyceraldehyde (3PGA), **k,** pyruvate **l,** lactate. TCA cycle intermediates **m,** citrate **n,** isocitrate **o,** acotinic acid **p,** alpha-ketoglutarate (aKG) **q,** succinate **r,** fumarate **s,** malate **t,** succinate/fumarate ratio. Intermediates of the polyol pathway **u,** sorbitol and **v,** fructose and intermediate of fructose metabolism **w,** fructose-1-phosphate (F1P) (n=4 for mouse, n=3 for naked mole-rat). Error bar represents mean ± s.e.m. n numbers refer to individual animals. Two-tailed, unpaired Student t-tests were used for statistical analysis in b-w. *p < 0.05, **p < 0.01, ***p < 0.001.

**Extended Data Fig. 6: Non-canonical glycogen pathways are altered in naked mole-rat heart**

**a,** expression of MGAM in heart determined by RNAseq (n=3), **b,** Fluorescence intensity quantification of GAA immunostaining of heart tissue sections in mouse and naked mole-rat at baseline and 30 mins ischaemia **c,** Mander's coefficient representing colocalization of GAA with glycogen in heart sections of mouse and naked mole-rat at baseline and 30 mins ischaemia (19-23 individual images were analysed, n=3-4). **m,** Representative images for **b** and **c**. Error bar represents mean ± s.e.m. n numbers refer to individual animals. Two-way ANOVA with Tukey's test was used for correction of multiple comparisons in **b,c,** *p < 0.05, **p < 0.01, ***p < 0.001.

Extended Data Fig.1: Metabolic response in ischaemic mouse and naked mole-rat heart and liver

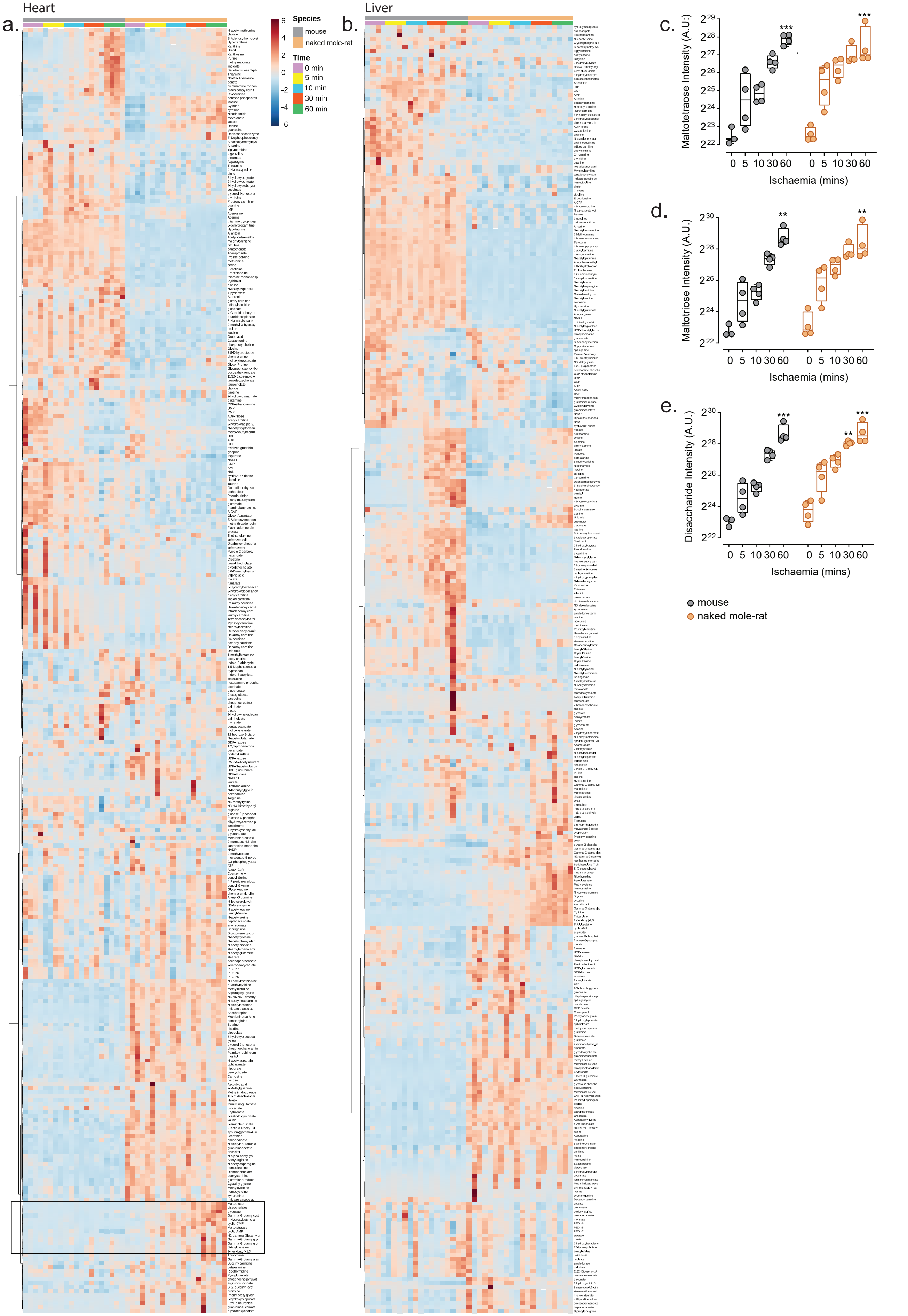

### Extended Data Fig 2: Expression of glycogen related genes in mouse and naked mole-rat liver

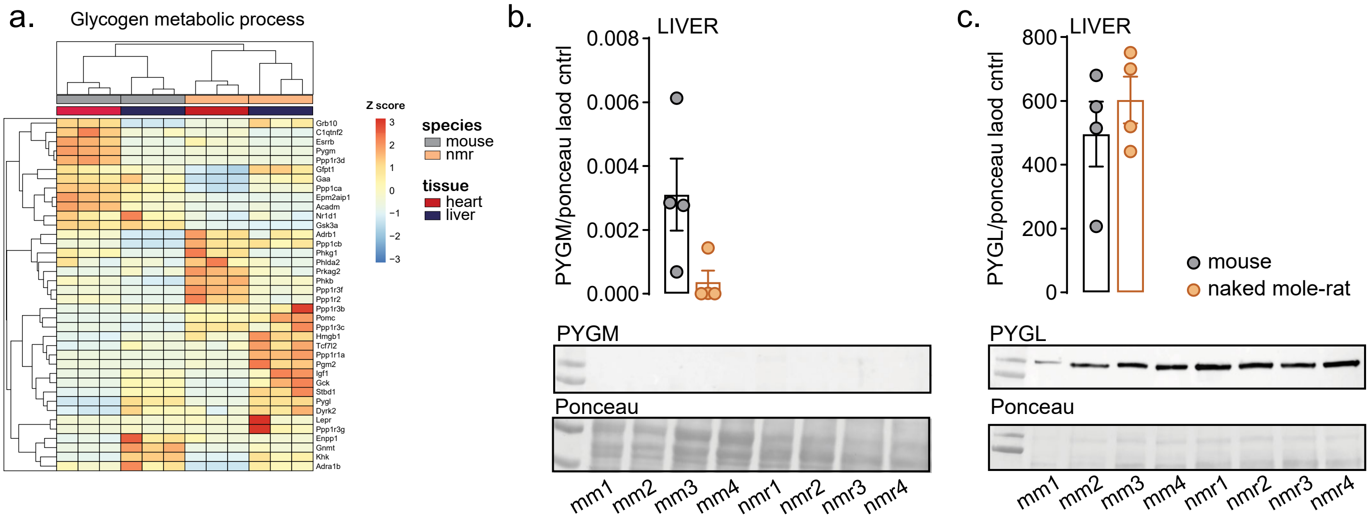

Extended Data Fig. 3: Glycolytic intermediates in ischaemic heart of mouse and naked mole-rat

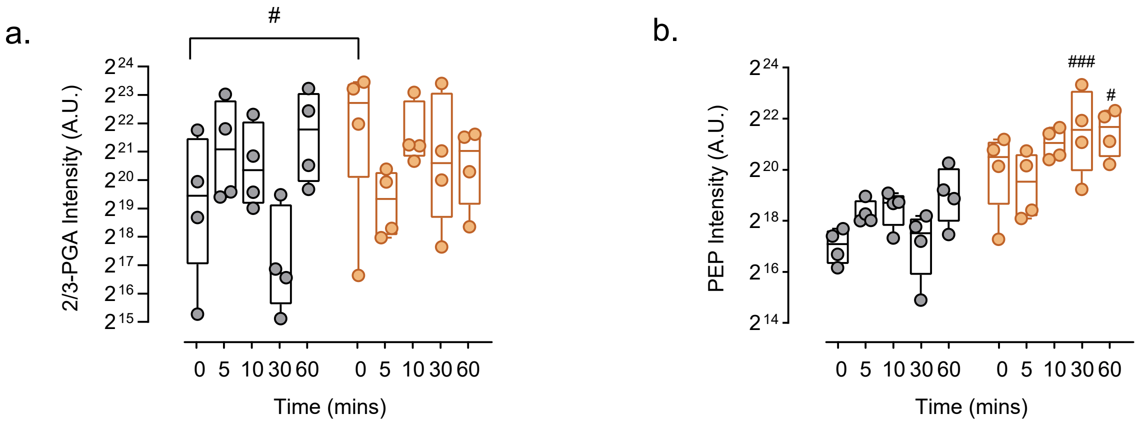

### Extended Data Fig.4: Co-option of Amy1 in naked mole-rat heart for efficient glycogen breakdown

a.

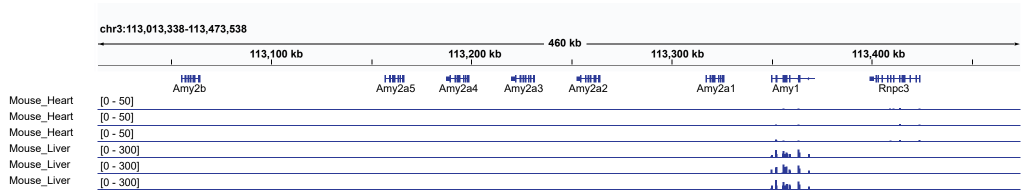


b.

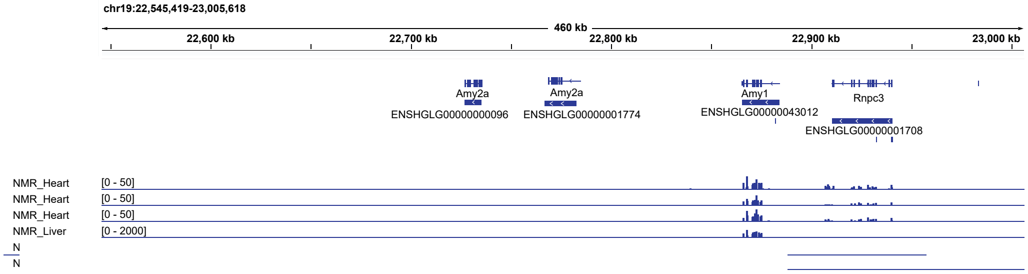


c.

| | HEART | | LIVER | | BRAIN | | |
|---|---|---|---|---|---|---|---|
| | nmr | mouse | nmr | mouse | nmr | mouse | |
| 55-65 kD | 24.0 | 21.8 | 26.8 | 26.0 | 22.5 | ND | ← expected MW of Amylase |
| 50-55 kD | ND | ND | 19.4 | 25.6 | ND | ND | |
| 45-50 kD | ND | ND | ND | ND | ND | ND | |
| 22-30 kD | ND | ND | ND | ND | ND | ND | |

Log₂(Intensity)

Extended Data Fig.5: Inhibition of amylase decreases glycolytic intermediates in hypoxia

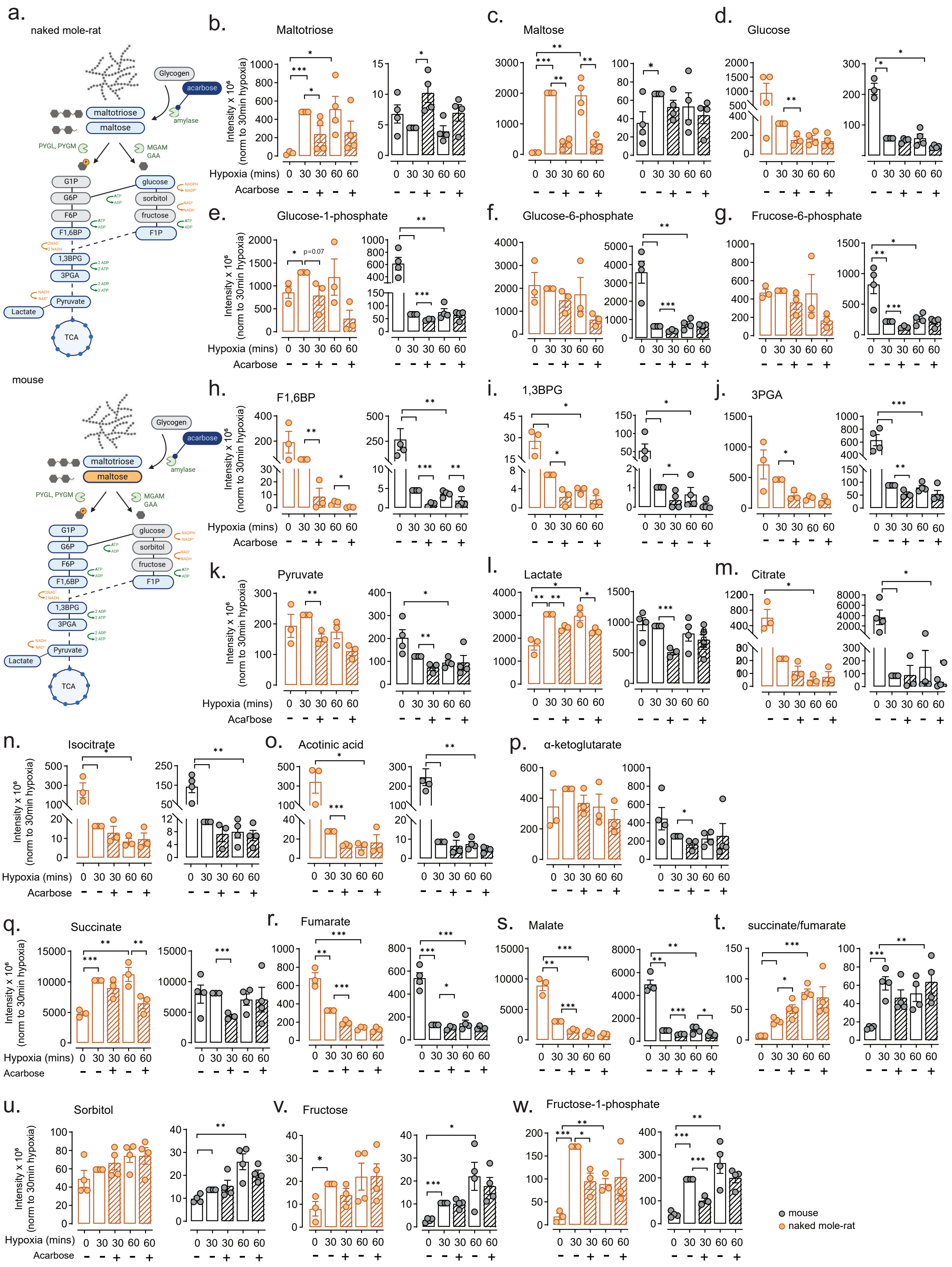

Extended data Fig.6: Non-canonical glycogen pathways are altered in naked mole-rat heart

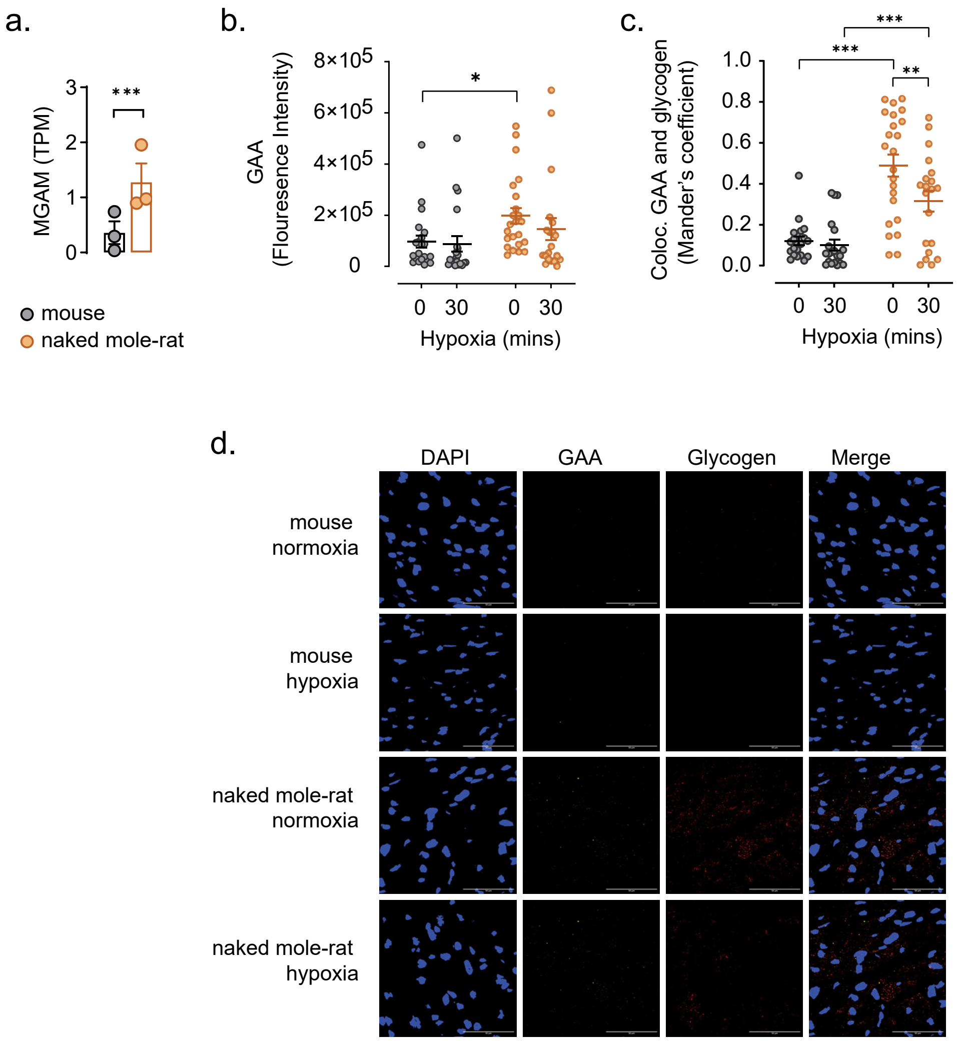
