## Supplemental Material and Methods for "Liver-like glycogen metabolism supports glycolysis in naked mole-rat heart during ischaemia"

### Materials and Methods

#### Animals

Adult male C57BL/6Nj mice (*Mus musculus*) were kept at 20-24°C in 12h light/dark cycles and fed ad libitum chow diet. Naked mole-rats (*Heterocephalus glaber*) were kept in individual colonies in a system of cages connected with plastic tubes within a humidified incubator (50–60% humidity, 28–30°C), in normal room air and darkness except during feeding and cleaning. Their diet consisted of fresh vegetables, fruit, and tubers (sweet potatoes) *ad libitum*. All water requirements were obtained from the food sources. Male and female non-breeding (subordinate) naked mole-rats were used in experiments since subordinate animals do not undergo sexual development or express sexual hormones[1]. The ages selected for this study allowed for physiological age matching such that all animals were at equivalent percentages of maximum lifespan and therefore, not the same chronological age. All animal procedures were conducted in accordance with European, national and institutional guidelines and protocols were approved by local government authorities Landesamt für Natur, Umwelt und Verbraucherschutz Nordrhein-Westfalen (LANUV).

#### Tissue sampling

For all experiments, adult mice were euthanized by cervical dislocation and adult naked mole-rats were anaesthetized in 4% isoflurane and euthanised by guillotine decapitation. Animals were not fasted before sampling. One day old neonatal mice (*Mus musculus*, C57BL/6Nj) were euthanised by decapitation. Tissues were quickly dissected out, snap-frozen in liquid nitrogen and stored at -80°C until use unless described otherwise.

#### Ischaemic tissue incubation

Adult male mice (12 weeks old, n=4, weight 23.2 g +/- 0.8 g, means +/- SEM) and adult naked mole rats (n=4, two male and two female, 2 years old, weight 33.6 g +/- 1.5 g, means +/- SEM) were euthanized as described above. The organs were removed and dissected as quickly as possible by three people in the order heart, brain, liver, kidney and quadriceps muscle and each tissue was cut into five portions of ~30-50 mg each. One piece was immediately frozen by freeze-clamping with Wollenberger tongs cooled in liquid nitrogen (time = 0 min). The remaining tissue pieces were incubated ischaemically in a humid environment at 32°C and sampled by freeze-clamping after 5, 10, 30 and 60 minutes. Tissues were stored at -80°C before metabolite extraction for LC-MS analysis.

In a separate experiment, one day old neonatal mouse pups (*mus musculus*, C57BL/6Nj, n=16) were killed by decapitation and the hearts and livers were either frozen immediately by freeze-clamping with Wollenberger tongs cooled in liquid nitrogen (time = 0 min) or incubated ischaemically in a humid environment at 32°C and sampled by freeze-clamping after 10, 30 and 60 minutes (n=4 each pr time point). Tissues were stored at -80°C before metabolite extraction for LC-MS analysis.

#### Hypoxic tissue incubation with amylase inhibitor

To detect the effect of amylase inhibition on ischaemic accumulation of polysaccharides and central carbon metabolism, adult male mice (12 weeks old, n=4, weight 22.5 g +/- 0.6 g) and adult naked mole-rats (n=4, two male and two female, 7-8 years old, weight 67.5 g +/- 3 g) were euthanized as described above. The heart was then quickly removed and the ventricle was cut into 6 pieces of ~50 mg. One piece was immediately frozen by freeze-clamping (time = 0).The remaining pieces were incubated in hypoxic ($pO_2 < 1\%$) modified Tyrode's buffer (10 mM HEPES, 6 mM KCl, 131 mM NaCl, 1 mM $MgCl_2$, 1.8 mM $CaCl_2$) with Vehicle or 2.5 mM acarbose (sigma A8980) dissolved in MilliQ Water, in an $O_2$ control glove box and cabinet (COY Laboratory) with oxygen concentration < 1%) for 15, 30 or 60 minutes before sampling by freeze-clamping and storage at -80°C for metabolomic analysis by GC and IC-MS and assaying for ATP concentration.

**Metabolite extraction for LC-MS metabolite analysis**

Frozen pieces of tissues (~10-20 mg) kept on dry ice were weighed and 40 µL/mg tissue extraction buffer (50% methanol, 30% acetonitrile and 20% ultrapure water with 5 µM valine-d8) was added on dry ice. Samples were homogenized on a Precellys 24 homogeniser at 4°C at 6500g 2x30s with 20s break. Metabolites were then extracted for 15 min at 4°C at 1500 rpm on an Eppendorf Thermomixer for 15 min before centrifugation at 4°C at 21000g for 20 min. The supernatant was transferred to glass autosampler vials and stored at -80°C until metabolite analysis by LC-MS.

**LC-MS data collection and analysis**

Metabolites were chromatographically separated using Millipore SeQuant ZIC-pHILIC analytical column (5 µm, 2.1 × 150 mm), coupled with a 2.1 × 20 mm guard column (both featuring 5 mm particle size). A binary solvent system was utilized, with solvent A comprising 20 mM ammonium carbonate and 0.05% ammonium hydroxide, and solvent B consisting of pure acetonitrile. The column oven temperature was maintained at 40 °C, while the autosampler tray was kept at 4 °C. Chromatographic elution was performed at a flow rate of 0.200 mL/min, following a programmed gradient: 0–2 min at 80% B, a linear decrease from 80% B to 20% B between 2-17 min, an immediate return to 80% B between 17-17.1 min, and a subsequent hold at 80% B from 17.1-23 min. Sample injection, each with a volume of 5 µl, was conducted in a randomized manner. Additionally, a pooled quality control (QC) sample, comprising equal proportions of all individual samples, was intermittently injected alongside the test samples.

A Vanquish Horizon UHPLC system coupled to an Orbitrap Exploris 240 mass spectrometer (both from Thermo Fisher Scientific) was employed to measure metabolite levels. The mass spectrometer was equipped with a HESI source, and the following parameters were used: spray voltages of +3.5kV/-2.8 kV, RF lens value of 70, heated capillary temperature of 320°C, and auxiliary gas heater temperature of 280°C. Sheath gas, aux gas, and sweep gas flow rates were set to 40, 15, and 0, respectively. MS1 scans were performed with a mass range of m/z=70-900, AGC target set to standard, and maximum injection time (IT) set to auto. Experimental samples were analyzed using full scan mode with polarity switching at an Orbitrap resolution of 120,000. Untargeted metabolite identification was performed using the AcquireX Deep Scan workflow, an iterative data-dependent acquisition (DDA) strategy with multiple injections of a pooled sample. The DDA full scan-ddMS2 method for the AcquireX workflow utilized the following parameters: full scan resolution of 60,000, fragmentation resolution of 30,000, fragmentation intensity threshold of 5.0e3, dynamic exclusion enabled after one time with an exclusion duration of 10s, mass tolerance of 5ppm, isolation window of 1.2 m/z, normalized HCD collision energies set to stepped mode (30, 50, 150), fragmentation scan range set to auto, AGC target set to standard, max IT set to auto, and mild trapping enabled.

Compound Discoverer software (v 3.2, Thermo Fisher Scientific) was used for metabolite identification. Metabolite identities were confirmed based on the following criteria: (1) precursor ion m/z matched within 5 ppm of the theoretical mass predicted by the chemical formula; (2) fragment ions matched within 5 ppm to an in-house spectral library of authentic compound standards analyzed with the same ddMS2 method, with a best match score exceeding 70; (3) metabolite retention times were within 5% of the retention time of a purified standard run with the same chromatographic method. Chromatogram review and peak area integration were performed using Tracefinder software (v 5.0, Thermo Fisher Scientific).

**LC-MS data analysis**

The generated table of metabolites intensities from peak integration with Trace Finder were processed to generate normalised Arbitary Units. Our R pipeline is based on the MetaProViz package (https://github.com/saezlab/MetaProViz) but we normalised the data using the probabilistic quotient approach[2,3]. In short, we filtered our data by Feature Filtering that applies the 80%-filtering rule on the metabolite features per condition[4], we impute missing values by half minimum imputation, performed per feature[4] and then we applied the PQN normalisation to account for unwanted variation. PCA plots generated with the prcomp function of r-cran stats package (https://search.r-project.org/R/refmans/stats/html/00Index.html) and heatmap plots were generated with the pheatmap function (https://CRAN.R-project.org/package=pheatmap.).

**Metabolite extraction for GC- and IC-MS metabolite analysis**

Frozen sections of heart tissue samples (~20-40 mg) were powdered for 1 min at 25 Hz on a TissueLyser (Qiagen) with pre-cooled metal balls, and metabolites extracted for 30 min at 4°C with shaking in 1 mL 40:40:20 (v:v:v) LC-MS Ultragrade acetonitrile, methanol and water with 20 ng/mL citric acid D4 internal standard. Protein was pelleted for 10 min at 21000g at 4°C and the supernatant was dried for 4-6h on a SpeedVac concentrator and stored at -80°C until analysis.

**GC-MS analysis of small molecules**

The analysis of polar metabolites was carried out using GC-MS (Gas Chromatography coupled to a Q-Exactive-Orbitrap mass spectrometer, Thermo Fisher Scientific). For this purpose, metabolites were derivatized using a two-step procedure starting with an methoxyamination (methoxyamine hydrochlorid, Sigma) followed by a trimethyl-silylation using N-Methyl-N-trimethylsilyl-trifluoracetamid (MSTFA, Macherey-Nagel). Analysis was performed as described previously [5] with slight modifications.

Dried samples were methoxyaminated by re-suspending them in 10 µL of a freshly prepared (40 mg/mL) solution of methoxyamine in pyridine (Sigma). The samples were incubated for 45 min at 40°C on an orbital shaker (VWR) at 1500 rpm. In the second step 90 µL of MSTFA spiked with C8 - C40 Alkane standard (40147-U, Sigma Aldrich) to a conzentration of 1 µg/ml was added and the samples were incubated for additional 45 min at 40°C and 1500 rpm. At the end of the derivatisation the samples were centrifuged for 2 min at 21100x g and the clear supernatant was transferred to fresh auto sampler vials with conical glass inserts (300 µl, Chromatographie Zubehoer Trott). For the GC-MS analysis 0.5 µL of each sample was injected using a TriPlus RSH autosampler system (Thermo Fisher Scientifc) using a Split/SplitLess (SSL) injector at 250°C in splitless mode. The carrier gas flow (helium) was set to 1ml/min using a 30m MEGA-5 MS capillary column (0.250 mm diameter and 0.25 µm film thickness, MEGA). The GC temperature program was: 1 min at 70°C, followed by a 9°C per min ramp to 350°C. At the end of the gradient the temperature is held for additional 5 min at 350°C. The transfer line and source temperature are both set to 280°C. The filament, which was operating at 70 eV, was switched on 4.5 min after the sample was injected. During the whole gradient period the MS was operated in full scan mode covering a mass range m/z 70 and 700 with a resolution of 60.000.

The GC-MS data analysis was performed using the open-source software El Maven [5] (Version 0.12.0). For this purpose, Thermo raw mass spectra files were converted to mzML format using MSConvert [6] (Version 3.0.22060, Proteowizard). The identity of each compound was validated by authentic reference compounds, which were measured at the beginning or at the end of the sequence; further by matching of the EI spectra and the retention index (RI).

For data analysis the peak areas of extracted ion chromatograms from selected fragment ions were determined with El Maven. The corresponding peak areas from isotopologue mass peaks of every required compound were extracted and integrated using the underlying algorithm within El Maven,. Extracted ion chromatograms were generated with a mass accuracy of <5 ppm and a retention time (RT) tolerance of <0.05 min as compared to the independently measured reference compounds. The pool size determination was carried out by summing up the peak areas of all detectable isotopologues per compound. These areas were then normalized to the tissue weight of each sample. For final analysis, samples were normalized by dividing normalized peak areas by its corresponding 30 minute hypoxia sample, thereby setting 30 minute hypoxia sample for each species to 1. In order to compare mouse and naked mole-rat, all samples were then rescaled by multiplying all hypoxia normalized peak areas by the average normalized peak area of the corresponding species at 30 minutes hypoxia.

**Anion-Exchange Chromatography Mass Spectrometry (AEX-MS) for the analysis of anionic metabolites**

Extracted metabolites were re-suspended in 150 µl of UPLC/MS grade water (Biosolve), of which 100 µl were transferred to polypropylene autosampler vials (Chromatography Accessories Trott, Germany) before AEX-MS analysis. The samples were analysed using a Dionex ionchromatography system (Integrion Thermo Fisher Scientific) as described previously[7]. In brief, 5 µL of the resuspended polar metabolite extract were injected in push-partial mode, using an overfill factor of 1, onto a Dionex IonPac AS11-HC column (2 mm × 250 mm, 4 µm particle size, Thermo Fisher Scientific) equipped with a Dionex IonPac AG11-HC guard column (2 mm × 50 mm, 4 µm, Thermo Fisher Scientific). The column temperature was held at 30°C, while the auto sampler temperature was set to 6°C. A potassium hydroxide gradient was generated using a potassium hydroxide cartridge (Eluent Generator, Thermo Scientific), which was supplied with deionized water (Milli-Q IQ 7000, Millipore). The metabolite separation was carried at a flow rate of 380 µL/min, applying the following gradient conditions: 0-3 min, 10 mM KOH; 3-12 min, 10−50 mM KOH; 12-19 min, 50-100 mM KOH; 19-22 min, 100 mM KOH, 22-23 min, 100-10 mM KOH. The column was re-equilibrated at 10 mM for 3 min.

For the analysis of metabolic pool sizes the eluting compounds were detected in negative ion mode using full scan measurements in the mass range m/z 77 – 770 on a Q-Exactive HF high resolution MS (Thermo Fisher Scientific). The heated electrospray ionization (ESI) source settings of the mass spectrometer were: Spray voltage 3.2 kV, capillary temperature was set to 300°C, sheath gas flow 50 AU, aux gas flow 20 AU at a temperature of 330°C and a sweep gas glow of 2 AU. The S-lens was set to a value of 60.

The IC-MS data analysis was performed using the open source software El Maven[6] (Version 0.12.0). For this purpose Thermo raw mass spectra files were converted to mzML format using MSConvert [8] (Version 3.0.22060, Proteowizard). The identity of each compound was validated by authentic reference compounds, which were measured at the beginning and the end of the sequence. For data analysis the area of the deprotonated $[M-H^+]^{-1}$ or doubly deprotonated $[M-2H]^{-2}$ isotopologues mass peaks of every required compound were extracted and integrated using a mass accuracy <5 ppm and a retention time (RT) tolerance of <0.05 min as compared to the independently measured reference compounds. Peak areas were then normalized tothe tissue weight of each sample.For final analysis, samples were normalized by dividing normalized peak areas by its corresponding 30 minute hypoxia sample, thereby setting 30 minute hypoxia sample for each species to 1. In order to compare mouse and naked mole-rat, all samples were then rescaled by multiplying all hypoxia normalized peak areas by the average normalized peak area of the corresponding species at 30 minutes hypoxia.

**RNA-seq sample preparation and data analysis**

Total RNA was isolated from three biological replicates per species and per group using RNeasy extraction kit (Promega). RNA-seq libraries were prepared using the Truseq stranded mRNA kit (Illumina) and sequenced using 100 bpPE read on the Illumina NovaSeq 6000 platform according to the manufacturer's instruction at Macrogen (Macrogen, Korea). Raw reads were filtered as adapter sequences and low-quality reads (-q 20) using Trim_Galore v0.6.10 (https://github.com/FelixKrueger/TrimGalore) via calling the Cutadapt[9] v3.6.dev2 tool. Reads were aligned to reference genomes GRCm39(mouse) and Naked_mole-rat_maternal(naked mole-rat) available on Ensembl database release110. For all species, the cleaned reads were aligned using Salmon[10] v1.10.0 to longest transcript of each gene extracted from corresponding genome annotations. For any gene with more than one transcript of same length, the transcript with the longest coding sequence was used. The orthologous transcripts between these two species were identified by performing reciprocal blast search (BLAST+ v2.10.0; blastp) against longest protein with parameters of "- evalue 1e-05; -max_target_seqs 1". The values of TPM, reads counts and effective gene length for each transcript were collected and integrated into transcript-sample table for each tissue according to all 15,512 identified 1:1 orthologous transcripts. The raw and normalized data are deposited at Gene Expression Omnibus (GEO, accession number GSE268095).

The counts tables were used to perform differential expression analysis between mouse and naked mole-rat with DESeq2 v1.42.0[11]. For each tissue count matrix, genes with sum reads counts <10 were filtered out. Since orthologous transcript lengths could vary between mouse and naked mole-rat genomes, we implemented an additional length normalization step in the DESeq2 pipeline to avoid biased comparative quantifications resulting from species-specific transcript length variation. To do this, the matrix of effective transcript lengths for each transcript in each sample was delivered to the DESeq2 'DESeqDataSet' object so that they are included in the normalization for downstream analysis. We considered all differentially expressed genes with an adjusted $p < 0.01$ (using the Benjamini-Hochberg algorithm) and $|\log_2(\text{fold change})| > 1.5$ to be statistically significant in our analysis. Weakly expressed genes, for which maximum detected expression level was > 1 TPM across all samples within each tissue were filtered out. Subsequently, we identified 1,512 up-regulated and 1,564 down-regulated genes in naked mole-rat heart in comparison to mouse, along with 2,159 up-regulated and 1,707 down-regulated genes in liver. Gene Ontology Analyses were performed using DAVID(https://david.ncifcrf.gov/summary.jsp)

**ATP assay**

Snap-frozen heart and liver tissues stored at -80°C for less than six months was assayed for ATP concentration by a bioluminescence-based luciferase assay (Strehler, 1974). Frozen tissue samples were homogenised on dry ice and extracted in perchloric acid (3% v/v HClO4, 2 mM $Na_2EDTA$, 0.5% Triton X-100) and diluted to 1 mg/mL. Samples and ATP standards were then pH neutralised with 2 M KOH, 2 mM $Na_2EDTA$ and 50 mM MOPS and the $KClO_4$ precipitate was pelleted by centrifugation at 10000g 4°C for 1 min. the supernatant was diluted 1:5 in buffer (100 mM Tris, 2 mM $Na_2EDTA$, 50 mM $MgCl_2$, pH 7.75). The [ATP] was measured by addition of luciferase/ luciferin solution (7.5 mM DTT, 0.4 mg/mL BSA, 1.92 µg luciferase/mL, 120 µM D-luciferin) to duplicates of samples and standards in a white 96-well microplate, incubated protected from light for 10 min before measuring bioluminescence using an Infinite 200Pro microplate reader (Tecan Technologies).

**Glycogen content assay**

Glycogen content of tissues were quantified using a Sigma glycogen assay kit (MAK 016) measuring absorbance at 570 nm on an Infinite 200Pro microplate reader (Tecan Technologies).

**TEM and confocal tissue sampling**

Hearts and livers from adult naked mole-rats (n=4, two female and two male, 2-7 years, 43.9 g +/- 10.1 g) and mouse (n=4, male, 8-12 weeks, C57BL/6Nj, 23.4 g +/- 1.2 g) were fixed for transmission electron microscopy and immunohistochemistry. One piece (~40 mg) from each tissue was immediately fixed in 4% formalin for IHC and one piece was cut into 1x1x1 mm pieces and immersion-fixed in 2% glutaraldehyde/ 2% formaldehyde (for liver in addition with 0.2% picrin acid) in 0.1 M cacodylate buffer (pH 7.2) overnight at 4°C. Another lump was incubated ischemically at 32°C in a humid atmosphere for 30 and 60 min before fixation in 4% formalin or cut into 1x1x1 mm pieces and immersion-fixed.

**TEM sample preparation and microscopy**
After fixation at 4°C overnight, the TEM tissue samples were rinsed in 0.1 M cacodylate buffer (pH 7.2) and incubated with 1% $OsO_4$ and 1% potassium ferrocyanid (liver samples) or in 2% $OsO_4$ (heart samples) in 0.1 M cacodylate buffer (pH 7.2) for 3 h (liver) or 2 h (heart) at 4°C. Subsequently, tissues were dehydrated using ascending ethanol series, transferred to propylene oxide and finally embedded in epoxyresin for 72 hours at 62°C. Ultrathin sections (70 nm) were cut with a diamond knife (Diatome, Biel, Switzerland) on an ultramicrotome (EM-UC6, Leica Microsystems) and placed on copper grids. Ultrathin sections were contrasted with 1.5% uranylacetate and lead citrate (Reynolds solution). Images were acquired with a transmission electron microscope (JEOL JEM 2100Plus), camera OneView 4K 16bit (Gatan), and software DigitalMicrograph (Gatan) at 80 kV at room temperature.

**TEM image analysis**
Lipid droplet density was quantified from 62 (mouse) and 52 (NMR) individual images of 3000 times magnification in Adobe Photoshop. A 1x1 µM grid was overlaid on the images and lipid droplet density was estimated by point-counting with grid intersections[12]. Glycogen granule diameters were measured from 25000 and 10000 times magnification images (7-12 individual images)and are expressed as mean feret diameter calculated by averaging the shortest and the longest measured segments of the granule using ImageJ.

**Immunohistochemistry GAA and glycogen**
After fixation for 24 hours at 4°C, formalin-fixed tissues were washed in PBS and stored at 4°C until dehydration and embedding in paraffin. The paraffin embedded tissues were sectioned on a Leica microtome at 4 µm thickness and placed on glass slides. Sections were then incubated at 60°C overnight and de-paraffinised and rehydrated in a series of xylol and ethanol washes, and antigens were exposed in citrate-based antigen unmasking solution (H-3300-250, Vector Laboratories) at 85-90°C for 10 min. The sections were then washed in PBS, permeabilized in 0.5% Triton-X-100, blocked in 5% BSA in PBS for 1h at RT and then incubated overnight at 4°C with anti-GAA (14367-1-AP, rabbit, Proteintech) and glycogen (mouse, kindly provided by Dr. Otto Baba). After washing in PBS, sections were incubated with secondary antibodies (goat anti-rabbit IgG (H+L) Alexa Fluor 488, A-11034, ThermoFischer and donkey anti-mouse IgG (H+L) Alexa Fluor 594, A-21203, ThermoFischer) for 1h at RT, then washed, stained with 300 µM DAPI (ab228549, abcam) and covered in ProLong Gold antifade mounting medium (Invitrogen). Sections were stored at 4°C in the dark until visualization.

Fluorescent signal was captured on a Airy Scan Confocal Microscope (Zeiss) with ×63 Plan-Apochromat/1.4 Oil DIC with ZEN software (Zeiss). Nuclei were visualized as DAPI staining at 358 nm, glycogen at 594 nm and GAA at 488 nm. Fluorescent signals were captured using the same intensity/recording settings for all sections. Fluorescent images were analysed using the 3D Suite and JACoP plugins in Fiji (ImageJ).

**Oil Red O**
Hearts from naked mole-rat and mouse were washed in PBS and frozen in Optimal cutting temperature compound (OCT compound). Then, tissues were sectioned on a Leica cryotome at 7 µm thickness and placed on glass slides. Slides were let to warm up at room temperature for 30 minutes. Slides were fixed for 10 minutes with 4% formaldehyde and washed in distilled water. Oil Red O (ab150678, abcam) staining kit was used following manufacturer's instructions. Slides were incubated in Propylene Glycol for 5 minutes and incubated overnight in Oil Red O solution, pre-heated at 60°C. Slides were incubated in 85% Propylene Glycol (diluted in distilled water) for 45 seconds, washed three times in water and mounted using Mounting Medium Aqueous (ab64230, abcam). Images were captured on a Slidescanner (S360, Hamamatsu) at x40. Images were analysed using the rawintden values from at least 30 randomly selected areas in Fiji (ImageJ).

**Amylase activity assay**

Snap-frozen tissues (~20 mg) stored at -80°C were homogenized in precellys tubes with zirconium beads in 1:10 weight:volume amylase buffer (7 mM NaCl, 50 mM KPi, pH 7.4) at 6500 g 2x30 sec on a Precellys 24 tissue homogenizer (Bertin Technologies). Homogenates were then centrifuged at 6500g for 10 min at 4°C and supernatants were mixed with 2.5 mM acarbose (dissolved in MilliQ water) or vehicle and assayed for total amylase activity on a Cobas 8000 modular analyzer (Roche Diagnostics).

**Western blotting**

Naked mole-rat and mouse heart and liver samples (n=4) stored at -80°C were homogenized in RIPA buffer (50 mM Tris, 150 mM NaCl, 1% NP-40, 10% SDS, pH 7.2 with cOmplete EDTA-free protease inhibitor tablets (Roche Diagnostics)) at 6500 g 2x30 sec on a Precellys 24 tissue homogenizer (Bertin Technologies). Homogenates were then centrifuged at 6500g for 10 min at 4°C, and the supernatants were aliquoted and stored at -80°C. Protein content was determined with a BCA assay using BSA as a standard. Samples were used on the first thaw, mixed 1:1 with 2x laemmli buffer and boiled at 95°C for 5 min to denature proteins. On a 10% acrylamide TGX Stainfree FastCast gel (Biorad), 30-60 µg protein/lane was loaded and the proteins were separated by electrophoresis for 20 min at 70V followed by 60-90 min at 120V until the loading dye reached the bottom of the gel. Proteins were then blotted onto nitrocellulose membranes by semi-dry transfer on a Biorad Trans-blot Semi-dry transfer cell at 25V for 15-30 min. Membranes were then blocked for 1h at RT in 3% BSA in TBST, and incubated with primary antibodies overnight at 4°C. Primary rabbit antibodies used were amylase (a8273, Sigma-Aldrich), PYGL (15851-1-AP, Proteintech), PYGM (15851-1-AP, Proteintech) and anti-GAA (14367-1-AP, Proteintech) using Histone H3 (ab1791, abcam) as loading control. After washing, membranes were incubated with rabbit secondary antibody (goat anti-rabbit HRP, A0545, Sigma) and proteins were visualized by chemiluminescence on a Chemidoc Gel Imaging System (Biorad) and total protein load was visualized with Ponceau S staining.

**Amylase qPCR and plasmid expression**

Total RNA was isolated from tissues using SV Total RNA Isolation System (Promega) in combination with TRI reagent (Sigma-Aldrich). After DNase digest, 1µg of total RNA was used for cDNA synthesis with iScript™ cDNA Synthesis Kit (Bio-Rad) following manufacturer's instructions. Measurement of cDNA level was performed with Luna Universal qPCR Master Mix (New England Biolabs) and the Bio-Rad CFX96 Real-Time PCR Detection System. The following primers were used to quantify Amy1 gene in mouse: 5′-GACTTTCCTGGAGTTCCCTATTC-3′ (forward), 5′-CGACAATCTCTGACCTGAGC-3′ (reverse) and naked mole-rat: 5′- GTTCCATACTCTGGTTGGGATT-3′ (forward), 5′-GACGACAATCTCTGACCTGAAA-3′ (reverse). To calculate absolute numbers of transcripts, plasmids were made containing the cDNA amplicon from each primer pair for mouse and naked mole-rat. The standard curve method with known doses of plasmid was used to quantitate mRNA transcripts by extrapolating a value by comparing unknowns to the standard curve of known transcript amounts.

**In-gel proteomics**

**Sample Preparation**

Mouse and naked mole-rat heart, liver and brain samples (n=1, ~20 mg each) were homogenized in RIPA buffer and proteins were separated by SDS-PAGE as described above on a 10% precast gel (Biorad) and fixed for 1h at RT in 10% acetic acid and 20% methanol in water. Bands of interests (55-65 kDa, 50-55 kDa, 45-50 kDa and 22-30 kDa) were excised and proteins were reduced and alkylated before digestion with trypsin before extraction of peptides with 30% acetonitrile / 3% trifluoroacetic acid in water. Samples were then acidified with 1% formic acid before loading onto SDB-RP StageTips before proteomics analysis.

**Data Acquisition**

Samples were analyzed by the CECAD Proteomics Facility on an Orbitrap Exploris 480 (Thermo Scientific, granted by the German Research Foundation under INST 216/1163-1 FUGG) mass spectrometer equipped with a FAIMSduo differential ion mobility device that was coupled to an Vanquish neo in trap-and-elute setup (Thermo Scientific). Samples were loaded onto a precolumn (Acclaim 5µm PepMap 300 µ Cartridge) with a flow of 60 µl/min before reverse-flushed onto an in-house packed analytical column (30 cm length, 75 µm inner diameter, filled with 2.7 µm Poroshell EC120 C18, Agilent). Peptides were chromatographically separated with an initial flow rate of 400 nL/min and the following gradient: initial 2% B (0.1% formic acid in 80 % acetonitrile), up to 6 % in 3 min. Then, flow was reduced to 300 nl/min followed by an increased of B to 20% in 26 min, up to 35% B within 15 min and up to 98% solvent B within 1.0 min while again increasing the flow to 400 nl/min, followed by column wash with 95% solvent B and reequilibration to initial condition. The FAIMS pro was switched between -40V and -60 V compensation voltages (CVs) with electrode temperatures kept at 99.5 °C for the inner and 85 °C for the outer electrode. The mass spectrometer was operated in data-dependent acquisition top 12 mode for both CVs with MS1 scans acquired from 350 m/z to 1400 m/z at 60k resolution and an AGC target of 300%. MS2 scans were acquired at 15 k resolution with a maximum injection time of 22 ms and an AGC target of 300% in a 1.4 Th window and a fixed first mass of 110 m/z. All MS1 scans were stored as profile, all MS2 scans as centroid.

**Sample Processing in MaxQuant**

All mass spectrometric raw data were processed with Maxquant (version 2.4) [13]using default parameters. with the match-between-runs option enabled between replicates. Samples stemming from mouse were analyzed against the canonical murine Uniprot reference proteome (downloaded 06/01/23). Naked mole rat samples were analyzed against an in-house built database which was created using Naked mole rat_maternal genome downloaded from Ensembl release110 and taking the longest protein sequence from each gene (see method for RNAseq analysis above). Gene Name is based on the orthologous transcript in mouse identified by performing reciprocal blast search between mouse and naked mole-rat (see method for RNAseq analysis above).For genes with missing annotated Gene Name, gene ID was indicated instead. Follow-up analysis was done in Perseus 1.6.15 [14]. Protein groups were filtered for potential contaminants and insecure identifications. Non-imputed, log2-scaled LFQ values were used for analysis. The mass spectrometry proteomics data have been deposited to the ProteomeXchange Consortium via the PRIDE[15] partner repository with the dataset identifier PXD052477.
